# Single-Molecule Tracking Reveals Dynamic Regulation of Ribosomal Scanning

**DOI:** 10.1101/2023.09.04.555162

**Authors:** Hea Jin Hong, Antonia L. Zhang, Adam B. Conn, Gregor Blaha, Seán E. O’Leary

**Author notes:** Eikon Therapeutics, Inc., 3929 Point Eden Way, Hayward, CA 94545. Bio-Rad Laboratories, Inc., 5731 W Las Positas Blvd., Pleasanton, CA 94588.

## Abstract

To initiate protein synthesis, the eukaryotic ribosomal pre-initiation complex must survey a messenger RNA leader sequence to identify the correct start codon.^1^ This pre-initiation complex motion through the leader, termed ‘scanning’, is coordinated by an intricate and highly-dynamic assemblage of translation factors, mRNA, initiator tRNA, and the small ribosomal subunit.^2,3,4^ Fundamental aspects of scanning dynamics remain poorly understood: estimates of its rate vary widely, and mechanisms that establish and regulate the motion remain largely unknown. Here we show, at the single-molecule level, that the *Saccharomyces cerevisiae* pre-initiation complex scans a diverse set of mRNA leaders at a rate of 10 – 20 nt s^−1^. Our data quantitatively support a scanning mechanism in which the mRNA leader is inspected base by base, essentially unidirectionally, and with modest sensitivity to mRNA structure. Unexpectedly, scanning bypasses canonical start sites where the initiator tRNA is present but GTP hydrolysis in the pre-initiation complex is impaired. Conversely, binding of the *S. cerevisiae* poly(A)-binding protein Pab1p to its own mRNA leader hinders scanning in a concentration-dependent manner. At saturating, physiological concentrations, Pab1p prolongs scanning by more than four-fold, evoking an autoregulation mechanism for translation initiation. Our data provide a real-time mechanistic framework for scanning regulation and energetics.

## Introduction

Faithful and efficient selection of the correct start codon is a critical aspect of protein synthesis. During eukaryotic translation initiation, a 43S pre-initiation complex (PIC) of the small (40S) ribosomal subunit, along with the methionylated initiator tRNA (Met-tRNA_i_^Met^) and protein initiation factors, is loaded at the mRNA 5’ end. The resulting 48S PIC locates a start site tens to hundreds of nucleotides distant. A ‘scanning’ mechanism for this search was proposed by Kozak nearly a half-century ago.^2^

In scanning, the PIC inspects the mRNA leader beginning near the 5’ end and proceeds in a 5’ to 3’ direction until it reaches an AUG start codon.^2,4,5^ The nucleotide context flanking this codon, conventionally termed the ‘Kozak sequence’, dictates the efficiency of its recognition; scanning that continues past the 5’-proximal start codon is termed ‘leaky scanning.’ ^4,6,7^

Recognition of, and commitment to an initiation codon is mediated through compositional and conformational changes in the PIC, which contains eukaryotic initiation factors (eIF)1, -1A, -3, and -5, and the GTPase eIF2 that delivers Met-tRNA_i_^Met^ to the ribosomal P site in a ternary complex with GTP. tRNA_i_^Met^ base pairing to an AUG triplet triggers rapid eIF1 ejection.^8^ eIF1 departure is accompanied by structural rearrangements, involving at least eIF1A and eIF5, that transition the initiator tRNA from a *P*_out_ to a *P*_in_ conformation with accommodation into the 40S P site.^9^ Closure of a latch on the 40S subunit then precludes further scanning.^10^ Start-site commitment is rendered irreversible through release of the phosphate product of eIF2 GTP γ-hydrolysis, gated by eIF1 ejection.^11^

While the steps underpinning start-site commitment are well understood, central aspects of scanning remain unclear. Though indirect bulk measurements have inferred a net forward scanning rate of ∼10 nt s^−1, 12,13^ a rate significantly over 100 nt s^−1^ was proposed in a recent single-molecule FRET study that probed the linear distance between the 48S PIC and an mRNA position 3’-proximal to the start site.^14^

Even earlier in initiation, the sequence of events that sets the stage for scanning is also incompletely understood. Bulk kinetic data from a reconstituted yeast system indicate a conformational change occurring shortly after PIC–mRNA recruitment limits the rate of start-codon recognition.^11^ Moreover, while early studies led to the proposal that arrival at a start codon leads to GTP hydrolysis, *in vitro* and *in vivo* data strongly suggest an equilibrium between intact and hydrolysed GTP is established on eIF2 prior to start-site arrival.^11,15^

The extent to which RNA *cis*-elements, beyond the Kozak sequence, might regulate scanning dynamics is also relatively unclear. For example, past data indicate significant differences in the scanning rate dependence on RNA structure.^14,16,17^ Genome-scale evidence captured by translation-complex profiling (TCP-Seq) indicates a complex relationship between scanning dynamics and leader identity.^18^ Emerging data also highlight an unexpected prevalence of initiation at codons near-cognate to AUG, i.e., differing by a single nucleotide.^19^

The extent to which RNA-binding proteins impede PIC progress through the leader is not known, though TCP-Seq data indicate such impedance may be significant. Protein modulation of scanning dynamics could constitute a *trans*-regulatory mechanism for initiation, and thus for translation.

We developed a single-molecule fluorescence assay to observe and quantify scanning dynamics on single mRNAs in real time with millisecond time resolution, with a reconstituted *S. cerevisiae* translation-initiation system. The assays were carried out in zero-mode waveguides (ZMWs), using a customized Pacific Biosciences *RS* II instrument,^20^ which permits simultaneous observation of hundreds of single-mRNA scanning reactions at physiological PIC and factor concentrations.

### Tracking single-mRNA scanning dynamics with eIF1

In our assay, eIF1 is leveraged as a reporter for the start and the end of scanning. Full-length, capped and polyadenylated mRNAs are surface-immobilised by poly(A)-tail capture, then pre-assembled 43S PICs, containing fluorescently-labeled 40S subunits, labeled eIF1, and other PIC components, are delivered to mRNA, to form 48S PICs (Figure 1a; Extended Data Figure 1a - c). PIC delivery occurs in the presence of the eIF4F cap-recognition complex (eIF4E•eIF4G1•eIF4A), eIF4B and ATP. 40S ribosomal subunits are fluorescently labeled with Cy3,^21^ eIF1 is Cy5-labeled at its C terminus,^22^ and mRNAs are Cy5.5-labeled *via* the immobilisation oligonucleotide.

**Figure 1.**
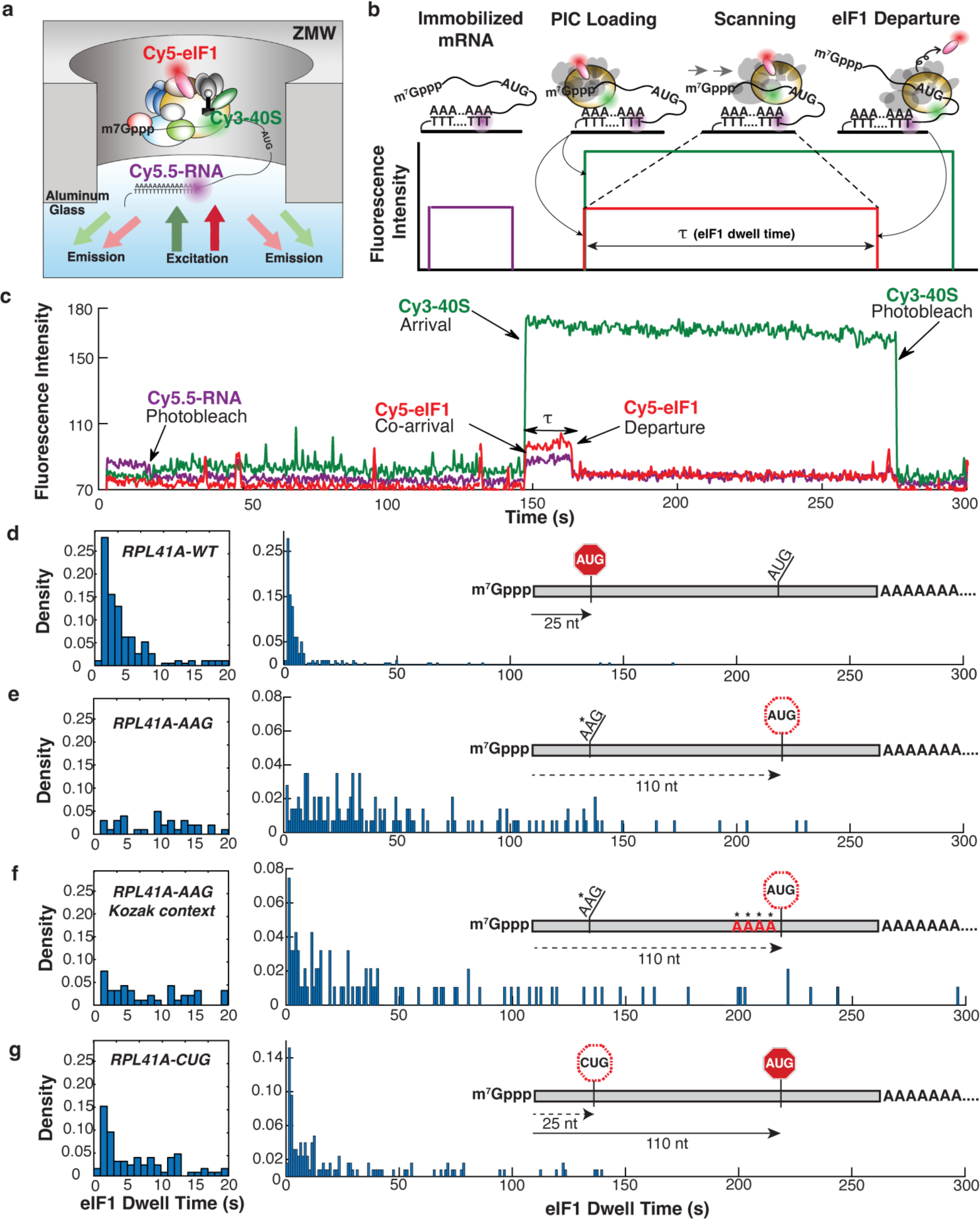
Direct observation of scanning in real time. **a.** Experimental setup for single-molecule fluorescence scanning assays in zero-mode waveguides. **b.** Idealised single-molecule fluorescence trace for scanning signal, indicating eIF1 dwell time, θ. **c.** Representative trace with scanning signal observed on the wild-type *RPL41A* mRNA. **d. – g.** Representative eIF1 dwell-time distributions on wild-type *RPL41A* mRNA (*n* = 193 molecules), *RPL41A*^AAG^ (*n* = 143), *RPL41A*^AAG/Kozak^ (*n* = 94) and *RPL41A*^CUG^ (*n* = 254) mRNAs, respectively. The distribution in the 0 to 20 s time domain is expanded to the left of the full-timescale distribution. Distributions from replicate experiments were statistically indistinguishable. Inset schematics indicate distances (in nucleotides) between the +1 nucleotide and the initiation codon.

At the single-molecule level, PIC–mRNA recruitment thus results in simultaneous appearance of Cy3-40S and Cy5-eIF1 fluorescence (Figure 1b). Both signals persist as the PIC scans, but the Cy5-eIF1 signal is lost upon start-codon recognition and eIF1 ejection. Thus, the eIF1 dwell time quantifies the time required for the PIC to prepare for scanning, scan the mRNA, recognize the start codon, and eject eIF1.

To validate that our system recapitulates on-pathway, cap-dependent translation initiation, we confirmed that labeled eIF1 is incorporated into PICs (Extended Data Figure 2a,b). eIF1 fluorescent labeling, *via* a fluorophore-conjugated C-terminal dipeptide extension, has been applied in bulk assays to quantify the eIF1 ejection rate upon start-codon recognition;^8^ a similar C-terminal extension was phenotypically silent *in vivo.*^22^ Efficient stable 40S subunit recruitment to capped mRNA also depended on eIF4F (Extended Data Figure 2c,d).^23, 24^

### A net forward scanning rate of ∼20 nt s^−1^

We applied our assay to characterise scanning on the yeast *RPL41A* mRNA, which has been extensively studied in bulk PIC-recruitment assays.^25,26,27^ We observed the expected Cy3-40S/Cy5-eIF1 co-arrival to immobilised *RPL41A* mRNAs, and subsequent loss of Cy5-eIF1 (Figure 1c).

The mean eIF1 dwell-time was 8.8 ± 1.9 s (mean ± standard deviation); the median time was 3.3 ± 0.1 s (Figure 1d). The dwell-time distribution was highly reproducible between replicate experiments (Extended Data Figure 3a,b). We expected the dwell time to be short on this 25-nucleotide leader, because its length is comparable to the 19-to 45-nucleotide 48S PIC footprint measured by TCP-Seq.^18^

The eIF1 dwell-time distribution rose to a short-time peak at ∼1 – 2 s that was followed by a quasi-exponential decay. The dominant exponential component implies a single kinetic step largely limits the PIC passage time from mRNA recruitment to eIF1 ejection on the *RPL41A* mRNA. Exponential fitting of the decay phase of the distribution (2 s ≤ τ ≤ 10 s) yielded a mean eIF1 dissociation rate of ∼0.36 ± 0.04 s^−1^, approaching the value of 0.6 s^−1^ for eIF1 ejection measured in bulk (Extended Data Figure 3c).^11^ The distribution was also well modeled by the probability density function of an irreversible 25-step process, with 24 steps occurring at a rate constant of 20 s^−1^ and a 25^th^ step occurring at 0.35 s^−1^ (Extended Data Figure 3d).^28^

To quantify the dependence of scanning time on leader length, we substituted the native *RPL41A* AUG start codon with AAG (*RPL41A*^AAG^). Although near-cognate to AUG, fungal PIC commitment to AAG initiation sites is low, at <0.5% efficiency relative to AUG;^29,30^ put otherwise, PICs typically scan past AAG triplets. This single U-to-A substitution caused the 5ʹ-proximal AUG codon to move 85 nucleotides downstream, to position +110.

eIF1 dwell times on *RPL41A*^AAG^ lengthened substantially (Figure 1e). The mean time was 59.6 ± 1.8 s, an almost seven-fold increase; the median time increased over eleven-fold, to 38 ± 4 s. If this increase reflected scanning to the +110 AUG triplet, it would imply a net rate of ∼2 nt s^−1^, much slower than expected based on past measurements. However, the *RPL41*^AAG^ +110 AUG triplet has a poor Kozak context (GATT<u>AUG</u>), raising the possibility it was not efficiently recognised. ^31^ Therefore, prolonged eIF1 dwell times may have resulted from bypass of the +110 AUG triplet and scanning to a downstream site. To test this possibility, we substituted the tetranucleotide sequence immediately upstream of the +110 AUG triplet with an optimal yeast Kozak sequence, AAAA<u>AUG</u>. Introduction of this sequence resulted in increased numbers of short eIF1 dwells, although the changes to the distribution did not reach statistical significance (Figure 1f). The median eIF1 dwell time was slightly shortened, to 34.0 ± 3.6 s, consistent with enhanced recognition of the +110 AUG codon. A long-dwell component also remained evident, as reflected in a slightly lengthened mean dwell time of 62.5 ± 3.8 s. Taken together, these data indicate the Kozak sequence was insufficient to promote full PIC commitment to the +110 AUG codon. Interestingly, of the RNA sequences studied in this work, the +110 *RPL41A* AUG is the only such triplet found in a 3’-untranslated mRNA region.

At the single-molecule level, multi-step processes where the observable signal remains unchanged, from the initial to the final step, are expected to produce a peaked, Erlang-type distribution for the time to complete all steps. Indeed, comparison of the short-time behaviour of the *RPL41A*^wt^ and *RPL41A*^AAG,Kozak^ eIF1 dwell-time distributions (Figure 1d, 1f) indicated a shift of the distribution to longer times for *RPL41A*^AAG,Kozak^ relative to *RPL41A*^wt^; this shift was more pronounced for *RPL41*^AAG^. If scanning were extremely rapid, it would not induce a delay on *RPL41A*^AAG^ or *RPL41A*^AAG,Kozak^ sufficiently long to cause this shift, as confirmed by simulation (Extended Data Figure 4). Therefore, the observed shift indicates the additional time scanning requires on the *RPL41A*^AAG^ mRNA approaches or exceeds that of the rate-limiting step that dominates the dwell-time distribution (1 / 0.35 s^−1^ = ∼2.9 s). Our data thus place the scanning rate on the order of 10 -20 nt s^−1^.

Recent *in-vivo* studies have highlighted the prevalence of initiation at near-cognate sites in yeast cells. Among near-cognate codons, CUG typically exhibits initiation efficiencies closest to AUG, though the estimated efficiency varies widely.^7,29,30^ Other near-cognate codons may be recognised with comparable efficiency in a suitable Kozak context.^32,33,34^ Indeed, in *in-vitro* bulk kinetic experiments with uncapped model mRNAs, AUG and even AUU triplets triggered eIF1 ejection with comparable efficiency.^8^ To quantify how a near-cognate triplet impacted scanning dynamics in the context of eIF4F-mediated initiation on a full-length, capped mRNA, we prepared an *RPL41A*^CUG^ variant, substituting CUG for the native (+25) start codon.

The mean and median eIF1 dwell times for *RPL41A*^CUG^ (28.9 ± 2.5 s and 9.2 ± 2 s, respectively) were between those for *RPL41A*^wt^ and *RPL41A*^AAG^. Interestingly, the dwell-time distribution on this *RPL41A*^CUG^ mRNA were markedly biphasic (Figure 1g; Extended Data Figure 3e,f), in contrast to *RPL41A*^wt^, *RPL41A*^AAG^, and *RPL41A*^AAG,Kozak^. Exponential fitting indicated the *RPL41A*^CUG^ distribution was an approximately 1:1 composite of those for *RPL41A*^wt^ and *RPL41A*^AAG^. Thus, our results are also consistent with *in-vivo* findings that the scanning PIC can efficiently recognise CUG start sites.

### Scanning dynamics on longer leaders

Our results for *RPL41A* raised the question of whether scanning dynamics would remain the same on longer leaders, as proposed from indirect bulk estimates of scanning rates.^12^ We thus measured the eIF1 dwell-time distribution on the 574-nucleotide *GCN4* leader, in a construct that includes the entire leader sequence and the first 30 nucleotides of the *GCN4* ORF. This leader contains four upstream open reading frames (uORFs), prior to the *GCN4* ORF. These uORFs leverage scanning dynamics to play a paradigmatic role in translational control.^35^ We serially substituted the uORF AUG triplets with AAG, producing five RNAs with varying cap-to-AUG spacing (Figure 2a).

**Figure 2.**
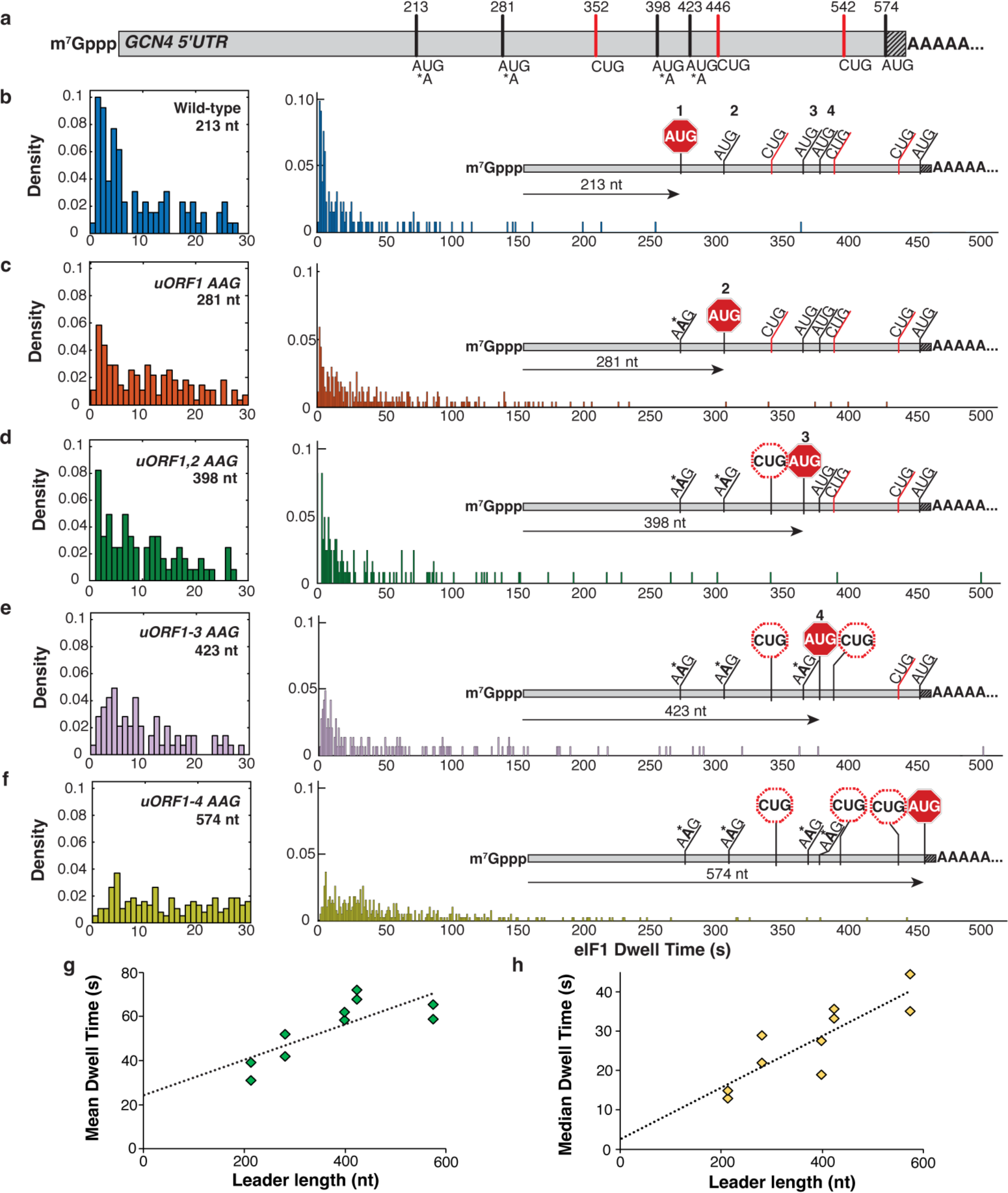
Scanning dynamics on the *GCN4* leader. **a.** Schematic of the *GCN4* leader indicating positions of canonical upstream ORF AUG codons, near-cognate CUG triplets, and the *GCN4* main-ORF AUG codon. The shaded portion indicates 30 nucleotides of the *GNC4* coding sequence included in the transcript. **b. – f.** eIF1 dwell time distributions for the wild-type *GCN4* leader sequence and the four sequential upstream ORF AUG-to-AAG variants. Left panels expand the short-time component of the distributions. Insets indicate relative linear distance traversed by the PIC for each leader sequence. For panels b. – f., the number of molecules in each distribution was 136, 115, 165, 98 and 160, respectively. **g. – h.** Mean and median eIF1 dwell times plotted against the cap to AUG leader length in nucleotides. Trendlines are from linear regression to all datapoints.

eIF1 dwell-time distributions systematically lengthened and changed in shape as the cap-to-AUG spacing increased. The distribution for the wild-type *GCN4* leader included a short-time peak at ∼2 s, followed by a quasi-exponential decay (Figure 2b). The mean and median dwell times for *GCN4*^wt^ were 35.1 ± 4.1 s and 13.9 ± 1.0 s. Since the wild-type *GCN4* leader includes 213 nucleotides between its 5’ end and 5’-proximal AUG, the observed distribution is again consistent with a net forward scanning rate of ∼10 nt s^−1^.

eIF1 mean dwell times increased to 47.0 ± 5.0 s, 60.1 ± 1.8 s, 69.9 ± 2.2 s, and 61 ± 3.3 s respectively for the serial uAUG *GCN4* variants; median dwell times also increased to 25.5 ± 3.5 s, 23.3 ± 4.3 s, 35.8 ± 2.5 s and 39.8 ± 4.7 s (Figure 2c–f). Linear regression of the dependence of these times on cap-to-AUG sequence length indicated increased eIF1 dwells of ∼0.08 s nt^−1^ (95% confidence intervals: 0.028, 0.13 s nt^−1^) and ∼0.065 s nt^−1^ (0.032, 0.099 s nt^−1^) based on analysis of the mean and median values, respectively (Figure 2g, h). These increased dwells correspond to scanning rates of ∼12 nt s^−1^ and ∼15 nt s^−1^, similar to our measurements for *RPL41A*. Thus, in their totality our data indicate that scanning rates of ∼12 – 20 nt s^−1^ are conserved across leaders that vary widely in length.

The *GCN4* leader additionally contains three CUG triplets, one located between the second and third upstream AUG codons, and two between the fourth upstream AUG codon and the *GCN4* ORF start codon (Figure 2a). While the linear relationship between eIF1 dwell time and cap-to-AUG sequence length indicates these CUG triplets are inefficiently recognised by the PIC, our data cannot rule out that some fraction of eIF1 ejection events result from their recognition. However, upstream CUG recognition would foreshorten the effective leader length. If that were the case, 12 – 15 nt s^−1^ would represent an upper limit on the true *GCN4* scanning rate.

As the effective leader length increased, the exponential component of the dwell-time distribution on *GCN4* was subsumed into a more symmetric distribution. This transition is consistent with PIC passage times becoming dominated by multi-step transit through the leader, rather than by a single kinetic step, again consistent with simulation (Extended Data Figure 4). However, the breadth of the observed distribution indicated that scanning rates differ substantially on individual molecules of the leader.

### AUG bypass with impaired GTP hydrolysis

Release of the phosphate product of eIF2-catalysed GTP hydrolysis occurs rapidly upon PIC commitment to a start site. While early mechanistic studies indicated that GTP hydrolysis is triggered by start-codon recognition,^36^ bulk pre-steady-state kinetic analysis in a reconstituted yeast initiation system suggested GTP hydrolysis occurs prior to or during scanning, and phosphate release from eIF2, subsequent to eIF1 ejection, is the physical process that cements irreversible tRNA_i_^Met^ accommodation in the 40S P site.^8,11^ However, these kinetic experiments were carried out with short unstructured and uncapped RNAs.

To delineate the contributions of GTP hydrolysis to scanning that follows cap-/eIF4F-dependent binding of the 43S PIC to a full-length mRNA, we determined the eIF1 dwell-time distributions for PICs formed with an eIF2 ternary complex that includes the GTP analogue, GDPNP, which is effectively non-hydrolysable on the timescale of our experiment. We compared the distributions for the *RPL41A*^wt^ mRNA and the *GCN4*^1–4^ ^AAG^ quadruple AUG mutant, which have 5’-proximal AUG sites at +25 and +574 nt from the cap, respectively (Figure 3a, b).

**Figure 3.**
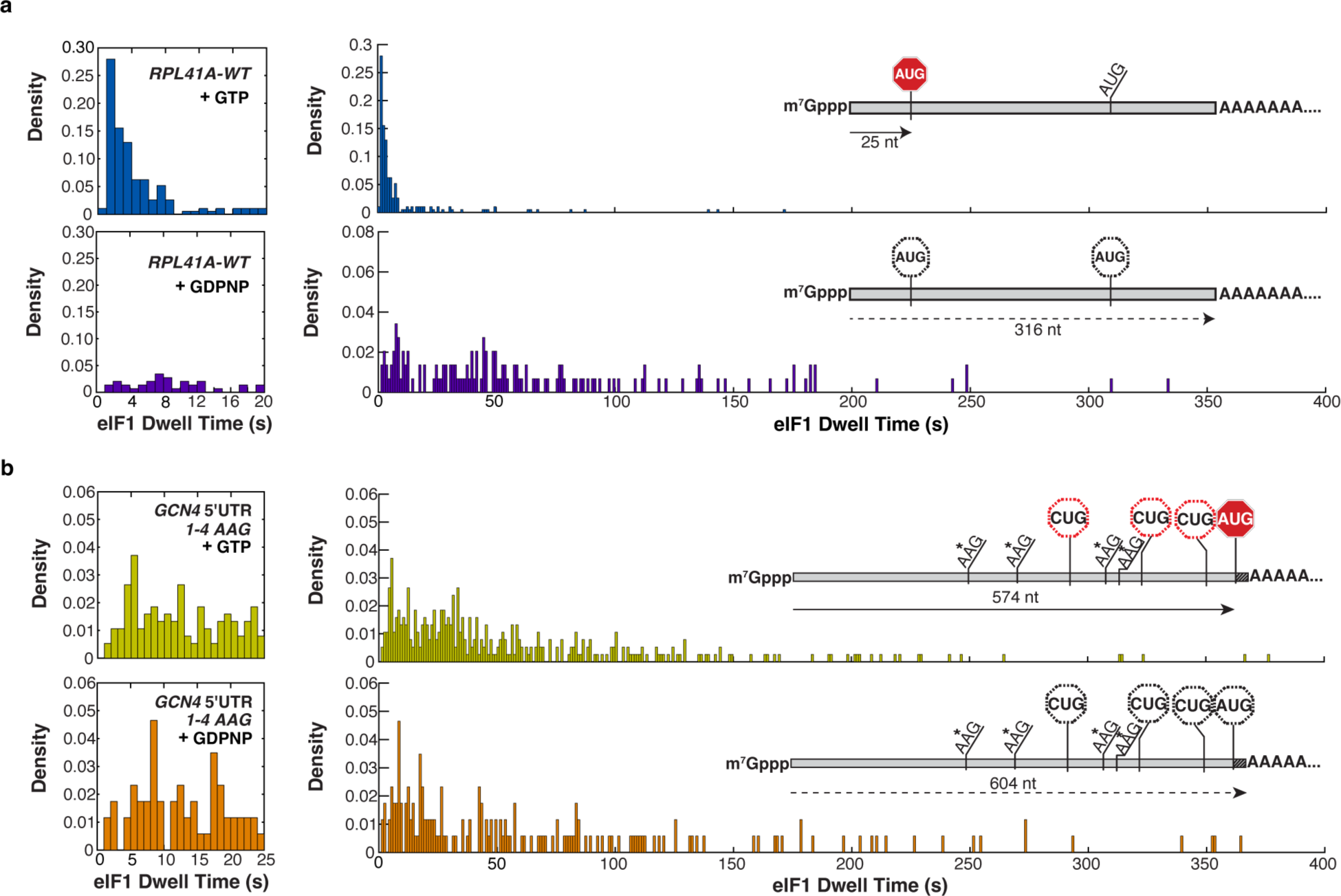
Regulation of scanning by GTP hydrolysis. **a-b.** eIF1 dwell-time distributions for scanning on (a) *RPL41A*^wt^ (*n* = 193 in the presence of GTP; *n* = 146 in the presence of GDPNP) and (b) the *GCN4* quadruple uORF AUG-to-AAG variant (*n* = 378 in the presence of GTP; *n* = 172 in the presence of GDPNP), contrasted between PICs formed with GTP *vs.* GDPNP in the eIF2 ternary complex.

The *RPL41A*^wt^ eIF1 dwell-time distribution was drastically altered in the presence of GDPNP. The mean increased about eight-fold, to 74 ± 5 s, while the median increased ∼15-fold, to 48 ± 1 s. The peak position increased from ∼1.4 s, to 5.8 s. This combination of changes is consistent with introduction of additional scanning steps, suggesting that GDPNP caused the PIC to bypass the +25 AUG start codon and continue moving along the mRNA.

If start-codon bypass were followed by continued scanning to the mRNA 3’ end, we hypothesised that inclusion of GDPNP in the ternary complex would have a lesser impact on eIF1 dwell times for *GCN4*^1–4^ ^AAG^. Because this RNA includes only ∼30 unhybridised nucleotides 3’ of its single AUG triplet, significant PIC motion past that triplet would be impossible. Indeed, the mean and median eIF1 dwell times were each lengthened for *GCN4*^1–4^ ^AAG^ by only ∼1.3-fold when GDPNP was included in the ternary complex (Figure 3a, b).

Our data are consistent with results from bulk PIC toeprinting assays with *RPL41A* that indicate the PIC is competent to scan in the presence of GDPNP. ^26^ However, in contrast to our results, the toeprinting assay detected PICs halted at the native *RPL41A* start codon when the ternary complex contained GDPNP, although eIF5 was absent. Conversely, our results mirror findings in a mammalian system that PICs bypassed AUG codons in poor Kozak context when GTP hydrolysis was impaired.^37^

Omission of eIF5, which serves as the GTPase-activating protein (GAP) for eIF2 within the PIC, also resulted in the shift of eIF1 dwell-time distribution parameters to longer times. Experiments with *GCN4*^wt^ and *PAB1*^AAG^ indicated both the short-time peak position and the mean and median dwell times increased, again consistent with scanning bypassing at least the first encountered AUG site when GTP hydrolysis is impaired (Extended Data Figure 5a,b).

### An RNA-binding protein blocks scanning

RNA-binding proteins have the potential to regulate translation efficiency *via* modulation of scanning dynamics, including by impeding passage of the scanning PIC through the leader. Indeed, TCP-Seq analysis for the *PAB1* mRNA, which encodes the poly(A)-binding protein Pab1p, indicated accumulation of 48S PICs at A-rich regions in its leader, including a sequence of 11 consecutive adenosines located +91 nt downstream of the cap. TCP-Seq and other data^18, 38^ have led to the proposal that Pab1p might negatively *trans-*regulate scanning dynamics on its own mRNA.

The eIF1 dwell-time distribution on the *PAB1* mRNA was biphasic, with a major component corresponding to rapid eIF1 ejection, similar to *RPL41A*^wt^, and a longer-time component with a mean at ∼17 ± 0.2 s and median of 4.9 ± 0.3 s (Figure 4a). The rapid-ejection population might suggest that scanning is intrinsically faster on *PAB1* than on *RPL41A* or *GCN4*. However, the *PAB1* leader includes two upstream near-cognate (CUG) initiation codons at positions +47 and +67 relative to the cap; the +67 CUG triplet is in a relatively good Kozak context (ATAT<u>AUG</u>). When these upstream CUG triplets were both substituted by AAG, the eIF1 dwell-time distribution became prominently peaked at ∼6 s (Figure 4b), consistent with introduction of additional scanning steps. The mean dwell time also increased (52 ± 5 s). Analysis of the distributions suggested the *PAB1*^wt^ eIF1 dwell-time behaviour was a composite of two parallel processes, the slower of which is common to *PAB1*^wt^ and *PAB1*^AAG^, and the faster of which is absent upon upstream CUG substitution. These results are consistent with the *PAB1*^wt^ eIF1 dwell-time distribution being generated by a composite of upstream CUG and *PAB1* ORF start-codon recognition.

**Figure 4.**
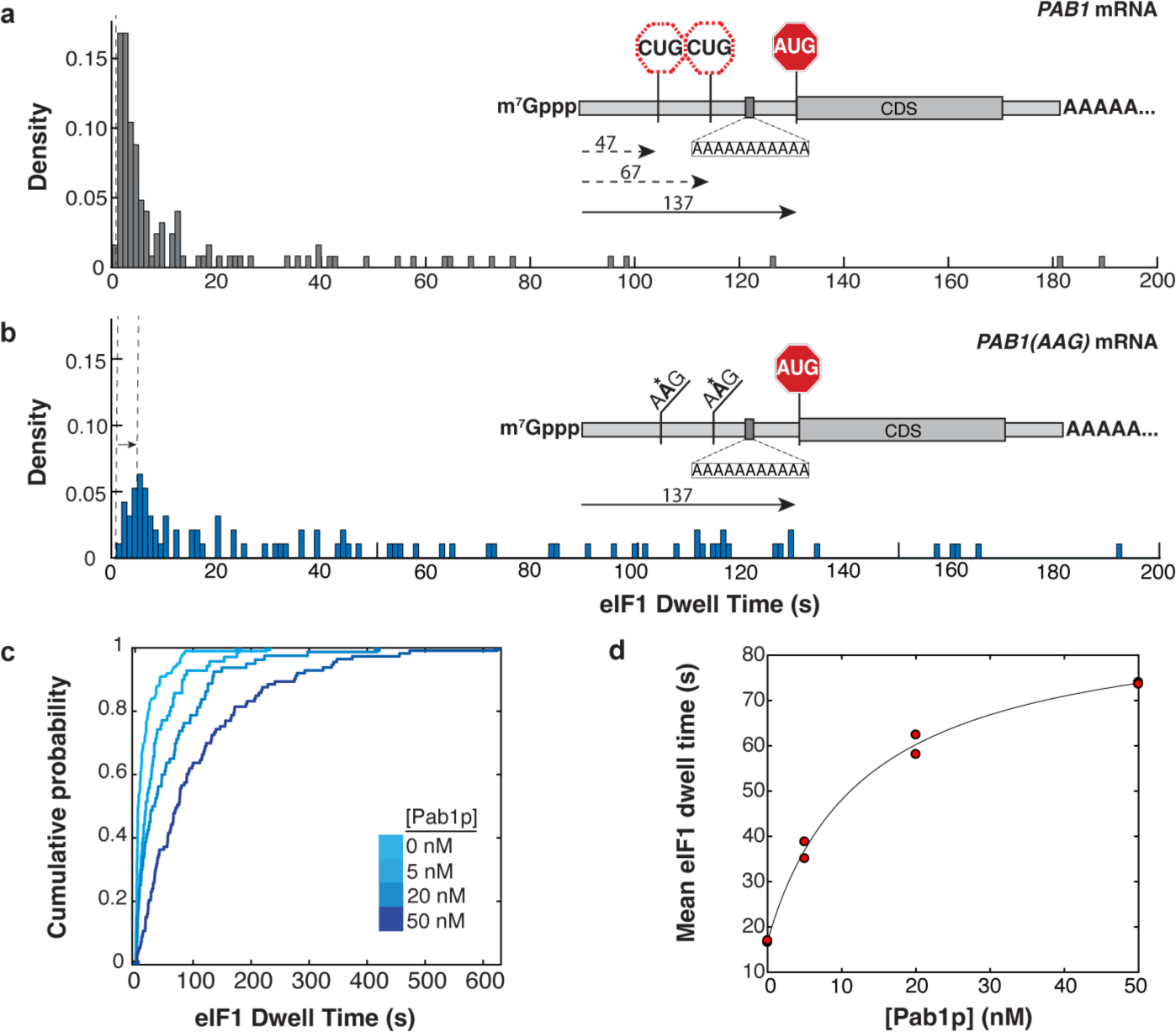
Regulation of scanning by *S. cerevisiae* poly(A)-binding protein. **a.** eIF1 dwell-time distribution for the *PAB1* mRNA (*n* = 125 molecules). Inset – relative locations of upstream CUG triplets and *PAB1* main-ORF AUG codon, and location of internal oligo(A) 11-mer. **b.** eIF1 dwell-time distribution for the *PAB1*^AAG^ mRNA, in which both upstream CUG triplets are substituted by AAG (*n* = 95). **c.** eIF1 dwell-time cumulative distribution functions for scanning on the *PAB1* mRNA in the presence of varying concentrations of the poly(A)-binding protein, Pab1p (5 nM, *n* = 110; 20 nM, *n* = 103; 50 nM, *n* = 112). **d.** Dependence of mean eIF1 dwell time on *PAB1* mRNA on the concentration of Pab1p, with hyperbolic fit (*K*_d_ 13 nM; 95 % confidence intervals 1.1, 25.6 nM). Replicate data points at 0 nM and 50 nM Pab1p overlap. The time added to scanning at saturating Pab1p concentration is 72 s (95 % confidence intervals: 64, 79 s).

The data indicate eIF1 ejection at the upstream CUG sites on the *PAB1* leader is significant, similar to the *RPL41*^CUG^ mRNA. Inspection of the TCP-seq profile for *PAB1* also reveals accumulation of 48S PICs in this region of the leader, at comparable abundance to accumulation at the *PAB1* ORF start codon.^18^ Interestingly, ribosome profiling suggests a low level of translation activity immediately downstream of these CUG codons, up to the position of an in-frame UAA stop codon at nucleotide +115 of the leader.^39^ However, the accumulation of PICs in this region of the leader is significantly greater than that of elongating ribosomes, based on comparison of the TCP-Seq and ribosome-profiling data.^18,39^ Evidently, a mechanism exists *in vivo* to suppress later stages of translation initiation, despite apparently high levels of recognition of this upstream initiation site.

To quantify the extent that Pab1p might impede PIC access to the *PAB1* start codon, we measured the eIF1 dwell-time distribution on the *PAB1*^wt^ mRNA in the presence of recombinant Pab1p. Addition of Pab1p shifted the distribution to longer times with increasing Pab1p concentration, in a saturable effect (Figure 4c). Moreover, when both upstream CUG triplets were mutated to AAG, the eIF1 dwell-time distribution shifted to a longer time and reduced the short-time component (Extended Data Figure 6a,b).

The mean eIF1 dwell time rose from 16.8 ± 0.2 s in the absence of Pab1p, to 37.0 ± 1.9 s, 60.3 ± 2.1 s, and 73.8 ± 0.2 s, at 5 nM, 20 nM, and 50 nM Pab1p. In contrast, addition of 50 nM Pab1p to the *RPL41A*^AAG^ mRNA, which lacks an internal poly(A) tract, slightly shortened eIF1 dwell times (Extended Data Figure 6c,d). This result strongly indicates the effects of Pab1p are directed by the *PAB1* internal oligo(A) sequence, rather than by interaction with the poly(A) tail or other PIC components such as eIF4G.

The eIF1 dwell-time dependence on Pab1p concentration for the *PAB1* mRNA was fit well by a hyperbolic function and the EC_50_ value from this fit (i.e., the concentration of Pab1p at which the eIF1 dwell-time prolongation was half-maximal) was ∼13 nM (Fig. 4d). Hill analysis of the data yielded similar results, and a Hill number (*n*_H_) of ∼0.98, consistent with Pab1p titrating a single site in the mRNA (Extended Data Figure 6e). The fitted EC_50_ value is close to the equilibrium dissociation constant of *S. cerevisiae* Pab1p from an oligo(A) 12mer, ∼30 nM;^40^ and also to the apparent equilibrium dissociation constant of Pab1p for full-length, non-polyadenylated *PAB1* mRNA (∼25 nM; Extended Data Figure 6f).

Taken together, our data indicate an mRNA-specific equilibrium-binding interaction of Pab1p with a single site in its own mRNA substantially impedes PIC from locating the *PAB1* start codon, but not the upstream CUG codons. These data are consistent with the impedance of scanning resulting from a steric block to PIC progress through the leader.

## Discussion

eIF1 dissociation from the PIC is a pivotal event in yeast translation initiation. Our real-time view of this process allows us to define the dynamics of scanning initiated by cap- and eIF4F-dependent PIC recruitment on full-length leader sequences.

Our measured scanning rate of ∼20 nt s^−1^ sets an upper limit of ∼16 kcal mol^−1^ on the average energy barrier for base-by-base PIC movement through the leader – assuming Eyring-type behaviour at each step. The true barrier height will be lower if scanning is not fully unidirectional.

Such a barrier is consistent with breakage of a few hydrogen bonds limiting PIC movement at each step (Figure 5). Since the tRNA_i_^Met^ anticodon includes three of the four RNA nucleotides, it can sample at least one base pair, or between two and five hydrogen bonds, at all non-AUG mRNA leader triplets except CCC. The stability of isolated tRNA duplexes with tri- or tetraribonucleotides lies in the region of -3 kcal mol^−1^;^41^ this includes an enthalpic contribution of around -15 kcal mol^−1^, which, in the spatially constrained environment of the PIC, is expected to dominate the barrier to tRNA-mRNA disengagement.^41^ Kinetic studies also indicate codon-anticodon base-pairing lifetimes in the 10 – 1000 ms range for cognate tRNA-tRNA pairs in solution, and ∼1 – 10 ms for near-cognate tRNA-tRNA pairs; again, these lifetimes might reasonably lengthen in the PIC.^42^ Thus, a scanning rate on the order of ∼10 – 100 nt s^−1^ is consistent with codon-anticodon base-pair dissociation limiting PIC movement during scanning.

**Figure 5.**
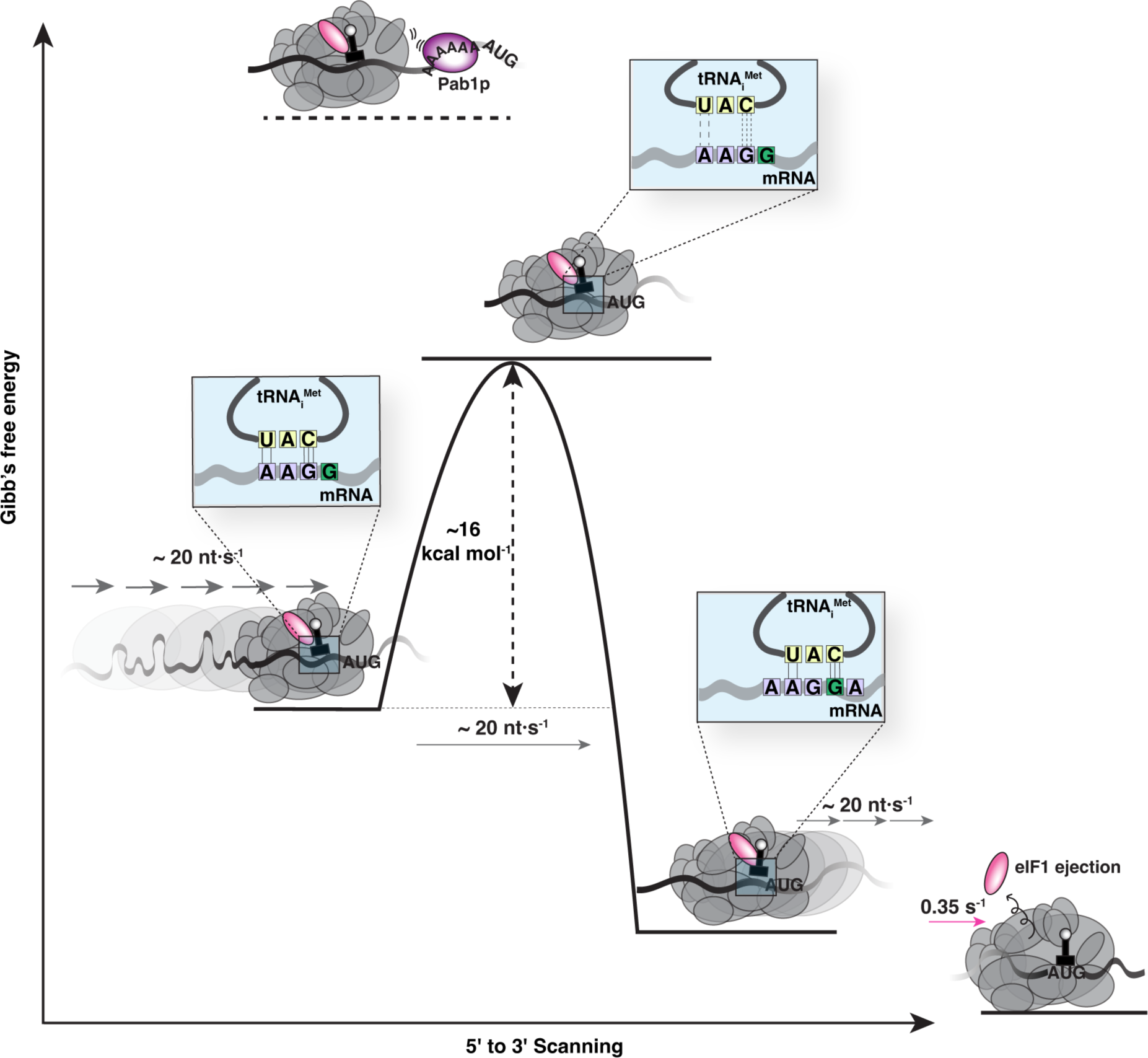
Energetics of scanning. The measured scanning rate of ∼20 nt s^−1^ implies an energy barrier of ∼16 kcal mol^−1^, on average, for PIC movement to the next nucleotide at each scanning step. The magnitude of this barrier corresponds to breakage of the hydrogen bonds in base-paired tRNA and mRNA; this base pairing by itself has an overall stability in the range of –3 kcal mol^-1^, with an enthalpic component of around –15 kcal mol^−1^. On the left of the schematic, one possible interaction during scanning includes five hydrogen bonds formed between a near-cognate mRNA triplet with a non-cognate tRNA. Thermal fluctuation that disengages the base pair is expected to traverse an activation barrier equal to or exceeding the enthalpy of the hydrogen-bond interactions, as measured by our assay. Steric hindrance from Pablp binding to an oligo(A) site abutting the PIC increases this activation barrier, resulting in blocked scanning. eIF1 ejection stabilizes the PIC to the point where no further forward motion is possible.

Considering the abundant evidence that mRNA-PIC interactions beyond the P site modulate scanning, e.g., the Kozak sequence, we propose these secondary interactions add to the total barrier height for each scanning step, tuning scanning rates. This is supported by structural data:^43^ the mRNA does not “float” over the PIC surface but is within van der Waals distance across a significant portion of the leader. Thus, a rate of 12 - 20 nt s^−1^ is quantitatively consistent with a scanning mechanism where the PIC actively surveys the leader by sampling tRNA-mRNA base pairing with a one-nucleotide step size.

Our *GCN4* data indicate scanning times vary linearly with leader length across a wide range. This is consistent with unidirectional or highly forward-biased motion. While we cannot rule out the possibility of low-amplitude local reversibility, as recently proposed,^44^ the observation that the eIF1 dwell-time dispersion does not significantly widen with leader length – e.g., *GCN4*^wt^ (213 nt) *vs. GCN4*^1–4^ ^AAG^ (574 nt) – supports past proposals of modest scanning reversibility.^12,13^

Nevertheless, we find wide variability in eIF1 dwell times between individual molecules of the same leader: scanning on a small proportion (4.6 %) of molecules for the 574-nt *GCN4*^1–4^ ^AAG^ leader is complete within 5 s, relative to mean and median times of ∼61 s and ∼40 s. We propose this variability results from mRNA structural heterogeneity that differentiates PIC passage times between leaders of identical sequence. Indeed, in a recent single-molecule FRET study, the signal ascribed to scanning became more heterogeneous at lower temperature, consistent with mRNA structure contributing appreciably to scanning dynamics.^14^

Our data reinforce the centrality of GTP hydrolysis by yeast eIF2 for start-site selection, but add to existing models with the finding that GTP hydrolysis is a prerequisite for eIF1 ejection. This contrasts with past observation of start-codon-dependent eIF1 dissociation from PICs programmed in a cap-independent manner with uncapped model RNAs.^8,11,26^ We suggest one or more steps of the PIC recruitment and scanning pathway on capped RNA enforce a GTP-hydrolysis requirement for eIF1 ejection. Consistent with this suggestion, eIF4F attenuates PIC GTP hydrolysis.^45^ Likewise, it has been suggested that addition of the mRNA cap and eIF4F enabled observation of eIF5 in a post-scanning PIC structure.^43^ The cap also blocks off-pathway PIC–mRNA recruitment.^26^ In sum, these past findings, and our data point to close communication between eIF4F and events occurring at the 48S P site during scanning.

Our results with GTPase impaired PIC also evoke the start-site bypass central to scanning-dependent translational control by upstream open reading frames. Conventionally this bypass is ascribed to PIC failure to acquire an intact ternary complex.^7^ We further find that, even when Met-tRNA_i_^Met^ can be delivered to the 48S P site, start-site bypass still occurs if GTP hydrolysis is impaired or blocked.

Taken together, our results point to a sustained driving force that ensures PIC directionality in scanning. A definitive answer to the mechanistic basis of directionality awaits further experimentation. However, our Pab1p/*PAB1* data directly calibrate its energetics. The hyperbolic dependence of mean scanning times on Pab1p concentration for the *PAB1* mRNA indicates that reversible Pab1p•*PAB1*-mRNA binding limits PIC passage through the leader. The extra passage time due to Pab1p-delayed scanning rises asymptotically with Pab1p concentration, to a maximum where all PAB1 leaders are blocked by Pab1p as the PIC reaches the Pab1p binding site. This is consistent with a sequential mechanism where the PIC arrives at the Pab1p blockage and is obliged to wait for Pab1p dissociation before further scanning (Extended Data Figure 7). The maximum delay time at saturating Pab1p concentration (72 s) indicates a Pab1p dissociation rate of ∼0.014 s^−1^, corresponding to a barrier height of ∼21 kcal mol^−1^. Evidently the PIC does not intrinsically have sufficient energy to move past a steric block of this magnitude, at least when the block is due to protein binding. Formation of the full complement of tRNA-mRNA hydrogen bonds upon P-site arrival of a cognate (AUG) start codon could impose a similar barrier, aiding the slowing of the PIC for eIF1 ejection and the structural changes that conclude scanning.

The finding that Pab1p significantly impedes PIC transit, specifically on its own *PAB1* mRNA, lends additional weight to past proposals that RNA-binding proteins may regulate initiation *in trans* through impacts on scanning dynamics. Further experiments will be required to assess the prevalence of this regulatory mechanism on a transcriptome-wide scale.

## Methods

### Purification of protein factors

Published constructs and strains were used to purify yeast eIF1,^23^ eIF1A,^23^ eIF5,^23^ eIF2,^46^ eIF3, eIF4E,^47^ eIF4A,^47^ eIF4G,^47^ eIF4B,^48^ Pab1p,^49^ 40S ribosomal subunit,^23^ tRNA,^23^ and the yeast methionyl-tRNA synthetase MetRS.^23^ All proteins were stored in 50 mM HEPES-KOH (pH 7.5), 100 mM KOAc, 3 mM MgOAc, 2.5 mM TCEP, 10 % glycerol except MetRS (40 mM Tris, 10 mM MgCl_2_, 10% glycerol, 2 mM DTT).

### Purification of eIF1/1A/5

Yeast eIF1, eIF1A, and eIF5 were purified as previously described.^23^ A pTYB2 vector containing the gene of interest, fused to a C-terminal intein-chitin-binding domain, was transformed into BL21 Codon Plus RIL cells (Agilent). The proteins were overexpressed with 1 mM IPTG, and the cells were lysed by sonication in lysis buffer (20 mM HEPES-KOH pH 7.4, 0.5 M KCl, 0.1 % TritonX-100, 1 mM EDTA, 1× protease inhibitor cocktail). The lysates were clarified and loaded on 1 mL chitin resin (NEB) and washed with 1 M KCl. The protein-bound resin was incubated overnight with cleavage buffer (20 mM HEPES-KOH pH 8.0, 500 mM KCl, and 75 mM DTT) to induce intein cleavage, then eluted the next day. For eIF1 C-terminal labeling, the protein was incubated with the cleavage buffer containing 200 mM 2-mercaptoethanesulfonate (MESNA) instead of DTT and Cys-Lys(ε-Cy5) overnight. The eluted proteins were further purified on a 1 mL heparin-HP column (Cytiva), followed by size-exclusion chromatography on Superdex 75. The purified proteins were flash-frozen with liquid nitrogen and stored at -80 °C.

### eIF2

The yeast strain, *GP3511*, which expresses N-terminally hexahistidine-tagged eIF2ϝ, in addition to eIF2α and eIF2β, was a generous gift from Dr. Alan G. Hinnebusch. The cells were grown in YPAD media (to a final OD_600_ of 4 – 6) and were harvested, washed, and resuspended (∼2 g of cells/1mL of lysis buffer; 30 mM HEPES-KOH pH 7.4, 800 mM KCl, 10 mM imidazole, 0.1 mM MgCl_2_, 5 mM β-mercaptoethanol, 10% glycerol, 1× protease inhibitor cocktail) and were flash-frozen with liquid nitrogen. The cells were lysed using a cryogenic mill (Spex Sample Prep, model 6875), for 15 repetitions of a two-minute run at 15 cycles per second, with 2 min cool time between repetitions. The debris was removed by filtration as previously described.^47^ The lysate was further clarified by centrifugation and then loaded onto a 5 mL Ni-NTA resin by gravity flow. The protein was eluted with lysis buffer containing 500 mM imidazole and was further purified using a heparin column followed by a Q column. The protein was then dialyzed into storage buffer. The pure protein was flash-frozen with liquid nitrogen and stored at -80 °C.

### eIF3

A yeast strain (*YOR361C*-TAP) bearing a genomically-encoded, C-terminally TAP-tagged eIF3b subunit, was purchased from Dharmacon. The growth condition was the same as for eIF2. The cells were harvested and washed with lysis buffer (10 mM Tris-HCl pH 8.0, 150 mM NaCl, 0.1 % NP-40, 1 mM PMSF). The washed cells were then resuspended in lysis buffer and flash-frozen with liquid nitrogen for storage until purification. The cells were lysed with a bead beater (45 s grinding followed by 2 min 15 s rest, for 5 cycles). The clarified and filtered lysate was loaded on 1 mL of IgG Sepharose fast flow resin (NEB). The column was washed with lysis buffer and TEV cleavage buffer (lysis buffer with 75 mM NaCl, 0.5 mM EDTA, and without DTT). The IgG-bound sample was cleaved with TEV protease overnight and further purified by size-exclusion chromatography on Superdex 200. The purified protein was flash-frozen with liquid nitrogen and stored at -80 °C.

### eIF4G

Yeast eIF4G was purified as described previously.^46^ A pTYB2 vector encoding a C-terminal fusion of eIF4G with an intein and chitin-binding domain was transformed into BL21 CodonPlus RIL cells (Agilent). Protein expression was induced with 0.5 mM IPTG for overnight at 16 °C. The cells were lysed by sonication in 50 mM HEPES-KOH pH 7.4, 500 mM KCl, 1 mM EDTA, 1 x protease inhibitor cocktail. The lysate was clarified before loading onto the chitin resin (NEB). The sample was treated with 3 U/μL of micrococcal nuclease (NEB) in 50 mM HEPES-KOH pH 7.4, 100 mM KCl, and 2 mM CaCl_2_ and was incubated with cleavage buffer containing 50 mM HEPES-KOH pH 7.4, 250 mM KCl, 1 mM EDTA and 50 mM DTT overnight. Eluted eIF4G was purified by anion-exchange chromatography on a 1mL Q-HP column (Cytiva) and dialyzed into the storage buffer, then flash-frozen with liquid nitrogen and stored at -80 °C.

### eIF4A/eIF4E

Yeast eIF4A and eIF4E were purified as described previously.^49^ A pET28a(+) vector encoding N-terminally his-tagged eIF4A or eIF4E was transformed into BL21 Codon Plus RIL cells. Protein overexpression was induced with 1 mM IPTG overnight at 16 °C. Cells were lysed by sonication, and the lysate was clarified by centrifugation before loading on the Ni-NTA column. For eIF4A, the protein eluted from the Ni-NTA step was further purified on a 1 mL Q-HP column followed by size exclusion chromatography (Superdex 75; Cytiva) in storage buffer. eIF4A was flash-frozen with liquid nitrogen and stored at -80 °C. For eIF4E, the Ni-NTA purified protein was buffer exchanged into the storage buffer over a 10-DG column (Bio-Rad) and was stored at 4 °C. Purified eIF4E preparations were used for no more than two weeks.

### eIF4B

A pET-22b(+) vector encoding hexahistidine-tagged eIF4B (*TIF3*) was transformed into BL21 CodonPlus RIL cells. The protein was purified as described previously.^50^ Protein overexpression was induced with 0.2 mM IPTG for 6 hours at 30 °C. The cells were resuspended in lysis buffer (50 mM Tris-HCl pH 7.5, 500 mM KCl, 20 mM imidazole) and lysed by sonication. The clarified lysate was loaded on 1 mL of Ni-NTA resin and was washed with lysis buffer containing 50 mM imidazole. The protein was eluted with lysis buffer containing 250 mM imidazole. The protein eluate was diluted into a low salt buffer (50 mM Tris-HCl pH 7.5, 100 mM KCl, 2.5 mM TCEP) and loaded on a pre-equilibrated 1 mL Q-HP column. The protein was further purified by size-exclusion chromatography (Superdex 200) in storage buffer. The purified protein was flash-frozen with liquid nitrogen and stored at -80 °C.

### Pab1p

A pET-28a(+) plasmid, encoding the *PAB1* gene fused to an N-terminal hexahistidine tag,^48^ was transformed into BL21-Codon Plus RIL cells. The cells were grown and lysed under the same condition as for eIF4E/4A, except the lysis buffer was composed of 50 mM HEPES-KOH pH 7.5, 300 mM KCl, 2.5 mM TCEP, 1× protease inhibitor cocktail, and 1× PMSF. The protein was loaded on 5 mL Ni-NTA resin and was washed with lysis buffer containing 10 mM imidazole. The resin-bound protein was treated with micrococcal nuclease, as for eIF4G. His_6_-Pab1p was then eluted with lysis buffer containing 250 mM imidazole. The protein was further purified on a 1 mL Q-HP column, and buffer exchanged into the storage buffer on a 10-DG column. The purified proteins were flash-frozen with liquid nitrogen and stored at -80 °C.

### 40S ribosomal subunit purification

Yeast 40S purification and labeling were carried out as described previously, with a few modifications.^21, 23^ The yeast strain encodes a 23-nucleotide extension to helix 39 of the rRNA which enables fluorescent labeling through hybridisation of a fluorophore-conjugated complementary oligonucleotide. Approximately 2 L of yeast culture was grown and harvested at O.D._600_ of ∼1 – 2. The cells were washed with lysis buffer (30 mM HEPES-KOH pH 7.5, 100 mM KCl, 20 mM MgCl_2_, 2 mM DTT, 2 mg/mL heparin, 20 U/mL RNAse inhibitor) and then flash frozen. The cells were lysed using a bead-beater and loaded on a 3 mL 1 M sucrose cushion. The sample was centrifuged in a 70Ti rotor at 55,000 rpm for 3.5 hours at 4 °C. The pellet was washed and resuspended with separation buffer (50 mM HEPES-KOH pH 7.5, 500 mM KCl, 2 mM MgCl_2_, 2 mM DTT). The resuspended sample was further clarified using a table-top centrifuge by centrifugation at 16,000 g for 15 min. 1 mM puromycin was added to the clarified sample, which was then incubated on ice for 15 min, followed by incubation at 37 °C for 10 min. The resulting sample was loaded onto a 5 – 20 % sucrose gradient, and then centrifuged for 10 hours at 22,000 rpm (SW-32 rotor). The gradient was pumped off with a peristaltic pump while monitoring the 254 nm absorbance and collecting 1 mL fractions. The fractions containing pure 40S subunits were pooled and buffer exchanged into storage buffer, then flash-frozen with liquid nitrogen and stored at -80 °C.

### *In vitro* transcription of mRNAs and purification

DNA templates for transcription of full-length mRNAs of *RPL41A* and *PAB1*, the *GCN4* leader, and a model AUG mRNA^23^ sequence were inserted into the pUC119 vector. A 5ʹ-GpG dinucleotide RNA sequence was added to each template sequence, to maximize transcription efficiency. A suitable restriction site was added to each template sequence 3ʹ end, to allow linearization for runoff transcription. Plasmids were transformed into DH5α (NEB). The transformed cells were grown in 300 mL of LB media at 37 °C overnight. Plasmid DNA was extracted using a maxi-prep kit (Takara Bio) and restriction-digested (*RPL41A* and *GCN4*-UTR templates with BamHI, *PAB1* template with EcoRI), to linearize the template at its 3ʹ end. The template was then transcribed with T7 polymerase (NEB), in transcription buffer (100 mM Tris-HCl, pH 8.0, 10 mM spermidine, 1 % TritonX-100, 16 mM NTPs, 3 % DMSO, 15 mM DTT, 25 mM MgCl_2_ and RNase inhibitor (NEB)). Reactions were incubated at 37 °C for 2 hours, followed by DNase treatment for 30 minutes. RNAs were extracted using phenol-chloroform-isoamyl alcohol (pH 4.3, 125:25:1) and ethanol-precipitated. The RNAs were capped with the *Vaccinia* capping enzyme (NEB) and poly(A)-tailed with *E. coli* poly(A) polymerase (NEB), following the manufacturer’s protocols. RNAs were freshly prepared before each single-molecule assay.

### Hybridization of mRNAs with Cy5.5-oligo

Capped and poly(A)-tailed mRNAs (1 µM) were hybridised to biotin-5’-d(T)_100_-3’-Cy5.5 (100 nM) in 100 mM HEPES-KOH pH 7.5 and 300 mM KCl, by heating to 98 °C for 3 min using a thermocycler with slow cooling to 4 °C at 0.1 °C s^−1^. The annealed product was used directly for single-molecule experiments.

### *In-vitro* transcription and purification of initiator tRNA

A pUC19 construct containing the yeast tRNA_i_^Met^ template sequence fused to a 3’ hammerhead ribozyme sequence was transformed into DH5α cells.^23^ tRNA_i_^Met^ was transcribed as for mRNAs, except on a larger (5 mL) scale for 4 hours, in 12 mM MgCl_2_. Ribozyme self-cleavage was initiated by increasing MgCl_2_ concentration to 30 mM and incubating at 60 °C for an additional hour. The tRNA was further purified by gel electrophoresis and gel extraction. The tRNA was buffer exchanged into 10 mM Bis-Tris-HCl pH 7.0 and 10 mM NaCl using a 5 kDa centrifugal filter (Milipore), and was stored at -80 °C.

### MetRS purification

A yeast strain harboring a plasmid that encodes GST-tagged methionyl-tRNA synthetase (MetRS) gene was grown in 3 L of YPAD media to ∼ 2 OD_600_. The cells were harvested and washed with 1× PBS, then processed as for purification of eIF3. The clarified lysate was loaded on a 1 mL GSTrap column (Cytiva), which was then washed with 1× PBS. MetRS was eluted with 10 mM reduced glutathione in 50 mM Tris-HCl pH 8.0 buffer. The eluted sample was buffer-exchanged into storage buffer (40 mM Tris-HCl pH 7.4, 10 mM MgCl_2_, 10% glycerol, and 2 mM DTT) on a 10-DG column (Bio-Rad). The purified protein was flash-frozen with liquid nitrogen and stored at -80 °C.

### Methionylation of Met-tRNA_i_^Met^ and acid-gel analysis

Yeast Met-tRNA_i_^Met^ was methionylated as previously described.^51^ The tRNA was refolded by heating at 70 °C for 10 min with 10 mM MgCl_2_ and slowly cooling to room temperature. The refolded tRNA was incubated with MetRS in acylation buffer containing 100 mM HEPES-KOH pH 7.6, 100 µM methionine, 10 mM ATP, 1 mM DTT, 10 mM KCl, and 20 mM MgCl_2_. The reaction was incubated at 37 °C for 20 min and extracted with acidic phenol, followed by chloroform-isoamyl alcohol (24:1). The methionylated tRNA was purified on Sephadex G-25. The acylated tRNA was analysed by acid urea-PAGE (acrylamide:bisacrylamide ratio of 19:1, 15 %, in 100 mM NaOAc pH 4.3) for verification and quantitation of the aminoacylation efficiency.

### Native PAGE analysis of PIC formation

PIC formation was verified by 4 % native PAGE as previously described.^23^ Ternary complex (TC) was formed by incubating eIF2 (800 nM) with GDPNP (2 mM) for 10 min at room temperature, followed by addition of Met-tRNA_i_^Met^ to 2 µM and incubation for another 5 min. For native-PAGE analysis, a 43-mer model mRNA (AUG) was used.^23^ The RNA was ligated to a 5ʹ-phosphorylated-Cy3-oligo at the 3’ end of the RNA by T4 RNA ligase (NEB) by following the manufacturer’s protocol. The Cy3-ligated AUG model mRNA was heated at 90 °C for 3 min and snap-cooled on ice for 5 min. The RNA was mixed with the TC, eIF1 (1 µM), eIF1A (1 µM), and 40S (100 nM) in the binding buffer (30 mM HEPES-KOH pH 7.5, 2 mM Mg(OAc)_2_, 100 mM KOAc, and 2 mM DTT). The mixture was incubated for an hour at room temperature and 10 % (w/v) sucrose was added before loading on the gel. The gel was electrophoresed for 1 hour 30 min at 4 °C and visualized using a Typhoon imager. This protocol was also used for analysis of PIC formation with Cy3-labeled 40S subunits.

### Pab1p electrophoretic mobility shift analysis

To quantify Pab1p binding to *PAB1* mRNA, the mRNA (50 nM) was incubated with varying concentrations of yeast Pab1p (200 nM to 1 nM) in 1× binding buffer (30 mM HEPES-KOH pH 7.4, 20 mM KCl, 1 mM MgCl_2_). The final salt and glycerol concentration of each sample was adjusted to 100 mM KCl and 10 % glycerol. The reaction was incubated for 30 min at room temperature, and then samples were loaded on a 0.8 % TBE (0.5×) agarose gel, pre-stained with ethidium bromide. The gel was electrophoresed at 80 V for 1.5 hours at 4 °C, and then visualized with a ChemiDoc imaging system (Bio-Rad).

### Single-molecule fluorescence assay

The scanning assay was performed in a customized PacBio RSII instrument.^51^ The general procedure described previously^49^ was modified as follows. For 40S subunit labeling, Cy3-oligo was incubated with 40S in ∼1.1:1 ratio (2.5 µM 40S : 2.25 µM Cy3-oligo), and the sample was heated at 42 °C for 2 min, then slowly cooled at 37 °C for 15 min and 30 °C for another 15 min. The ternary complex was formed by incubating eIF2 (2 µM) with GTP (3.5 mM) for 10 minutes, and then with initiator tRNA (2 µM) for a further five minutes, at room temperature. Initially, a SMRT cell (Pacific Biosciences) containing 150,000 ZMWs was reacted with 16 µM NeutrAvidin (Invitrogen) for 5 minutes at room temperature. The surface was then washed with assay buffer three times (30 mM HEPES-KOH pH 7.5, 3 mM Mg(OAc)_2_, and 100 mM KOAc). The biotin-5’-d(T)_100_-3’-Cy5.5 annealed mRNA sample (10 nM) was immobilized for 5 min. The surface was again washed with assay buffer, and then blocked with unlabeled eIF1 (1 µM), a mixture of Biolipidure 203 and 206 (5 % (v/v) each, NOF America), and 1 mg/mL BSA, to abrogate non-specific surface binding of Cy5-eIF1 and Cy3-40S subunits. Excess block-mixture components were removed by one wash with assay buffer supplemented with 2 mM protocatechuic acid (PCA), 2 mM triplet-state quencher (TSY; Pacific Biosciences), and 1× protocatechate-3,4-dioxygenase (PCD) oxygen scavenging system (Pacific Biosciences). 20 µL of imaging buffer containing the oxygen scavenging system and 1 mg/mL BSA was then added to the SMRT cell. During cell preparation, PICs were reconstituted by combining 50 nM Cy5-eIF1, 500 nM eIF1A, 50 nM Cy3-40S, 300 nM TC, 500 nM eIF3, 200 nM eIF4E, 80 nM eIF4G, 1 µM eIF4A, 500 nM eIF4B, (varying concentration (10 - 100 nM) of Pab1p was added to experiments containing Pab1p for *PAB1* mRNA), 2 mM ATP, 200 nM eIF5, oxygen scavenging system, and 1× casein, in the order given. The final concentration of all components is halved with respect to the values above, upon dilution of the sample into the 20 µL of imaging buffer previously added to the cell. The reconstituted PIC (20 µL) was then robotically delivered at the beginning of the experiment and Cy3, Cy5, and Cy5.5 fluorophores were excited by illuminating with green (532 nm) and red (642 nm) laser at 0.7 µW µm^-2^ and 0.07 µW µm^-2^ respectively. Movies were recorded at 10 frames per second for 15 min. For the experiments with GDPNP, the TC was formed with GDPNP instead of GTP.

### Single-molecule data analysis

The movies were analyzed using MATLAB scripts reported previously.^51^ Fluorescence traces were first extracted from raw movie data with automated selection of ZMWs containing a Cy3 signal. The resulting traces were then manually curated to ensure they each contained: 1) a Cy5.5-mRNA signal at the start of the movie, and 2) an event with co-arrival of Cy3-40S and Cy5-eIF1. Put otherwise, traces were only included in the dataset if they demonstrably reported on a surface-immobilised RNA being recognised by a complex of eIF1 with the 40S subunit at the ZMW surface. The durations of the Cy5-eIF1 events that followed co-arrival with a Cy3-40S signal were manually assigned and tabulated. Data were binned in 1-second increments to produce the normalized-density eIF1 dwell-time histograms depicted in the Figures.

### Statistical methods

All statistical analysis and fitting was performed in MATLAB. Wilcoxon rank-sum tests to compare dwell-time distribution pairs were performed at a significance level of 0.05. For empirical exponential fitting of eIF1 dwell-time distributions, the tabulated dwell times were converted to an empirical cumulative distribution function, then this function was fit by non-linear regression using the Trust-Region algorithm with bisquares weighting. Goodness-of-fit was evaluated by inspection of root-mean-square error values, which were typically in the range of probability-0.01 to -0.09. An identical non-linear regression approach was taken for hyperbolic and Hill fitting of the scanning time dependence on Pab1p concentration. For linear regression of mean or median eIF1 dwell times versus GCN4 leader lengths, the data were fit using the Trust-Region algorithm. Goodness-of-fit was evaluated based on the R-square value, which ranged from 0.7 to 0.8. 95% confidence intervals for estimation of the slopes are reported in the text.

## Acknowledgements

We thank Alan G. Hinnebusch (National Institute of Child Health and Human Development) for providing yeast strain *GP3511*, and Joseph D. Puglisi (Stanford University School of Medicine) for providing the strain for production of the fluorescently-labelable yeast 40S subunits. pUC19-HHtRNA, for preparation of the initiator tRNA, was a gift from Jon Lorsch (AddGene plasmid #37222). Plasmid pSW149, for eIF4G1 overexpression, was a gift from Sarah Walker (University at Buffalo).

## Author contributions

S.O’L conceived the study. H.H. and A.Z. performed the experiments. A.Z., and H.H. purified and labeled the ribosomal subunits, with contributions from A.C.. H.H., A.Z., G.B., and S.O’L analyzed the data. S.O’L, and H.H. wrote the manuscript, with contributions from G.B., and in consultation with all other authors

## Data availability

All data acquired and analysed in this study are included in the manuscript and Extended Data.

## Code availability

Custom code, implemented in MATLAB, for extraction and analysis of single-molecule fluorescence trajectories is available at https://drive.google.com/drive/folders/1k1ZvZqb-TMgjpysfUVnEEwbiFJ7s0rsr?usp=sharing

Correspondence and requests for materials should be addressed to Seán O’Leary.

---

**Extended Data Figure 1.**
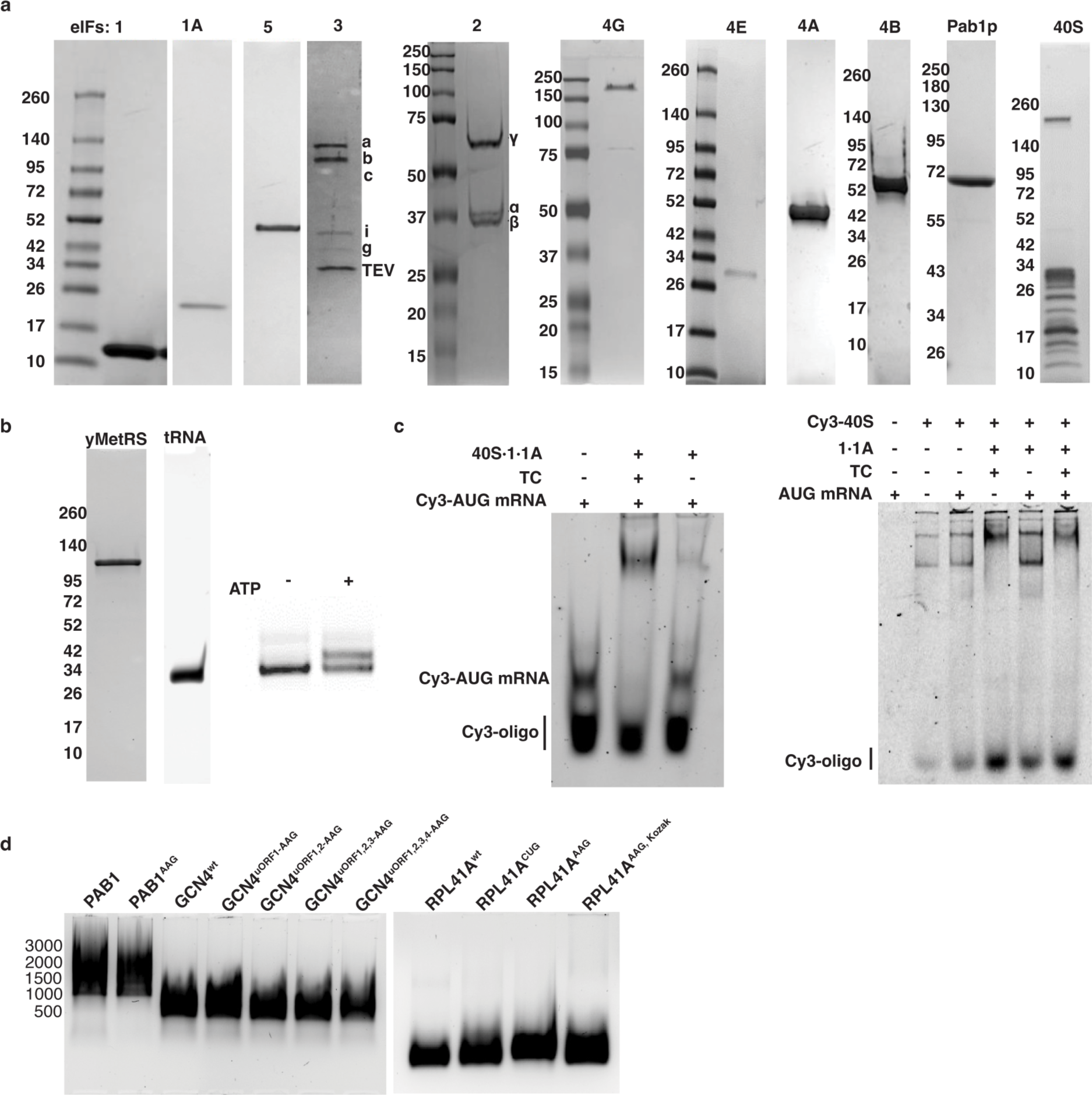
Purification and validation of reagents. **a.** SDS-PAGE analysis of purified individual components of the yeast PIC. **b.** SDS-PAGE analysis of yeast methionyl transferase (yMetRS) and urea-PAGE analysis of *in vitro* transcribed tRNA. Acid-gel analysis of methionylation efficiency (right) **c.** EMSA of the reconstituted PIC containing Cy3-model (*AUG*) mRNA, eIF1, eIF1A, and the eIF2 ternary complex formed with GDPNP (left). The gel on the right represents PIC formation with the Cy3-labelled 40S subunit. **d.** *In vitro* transcribed RNAs on a TBE agarose gel.

**Extended Data Figure 2.**
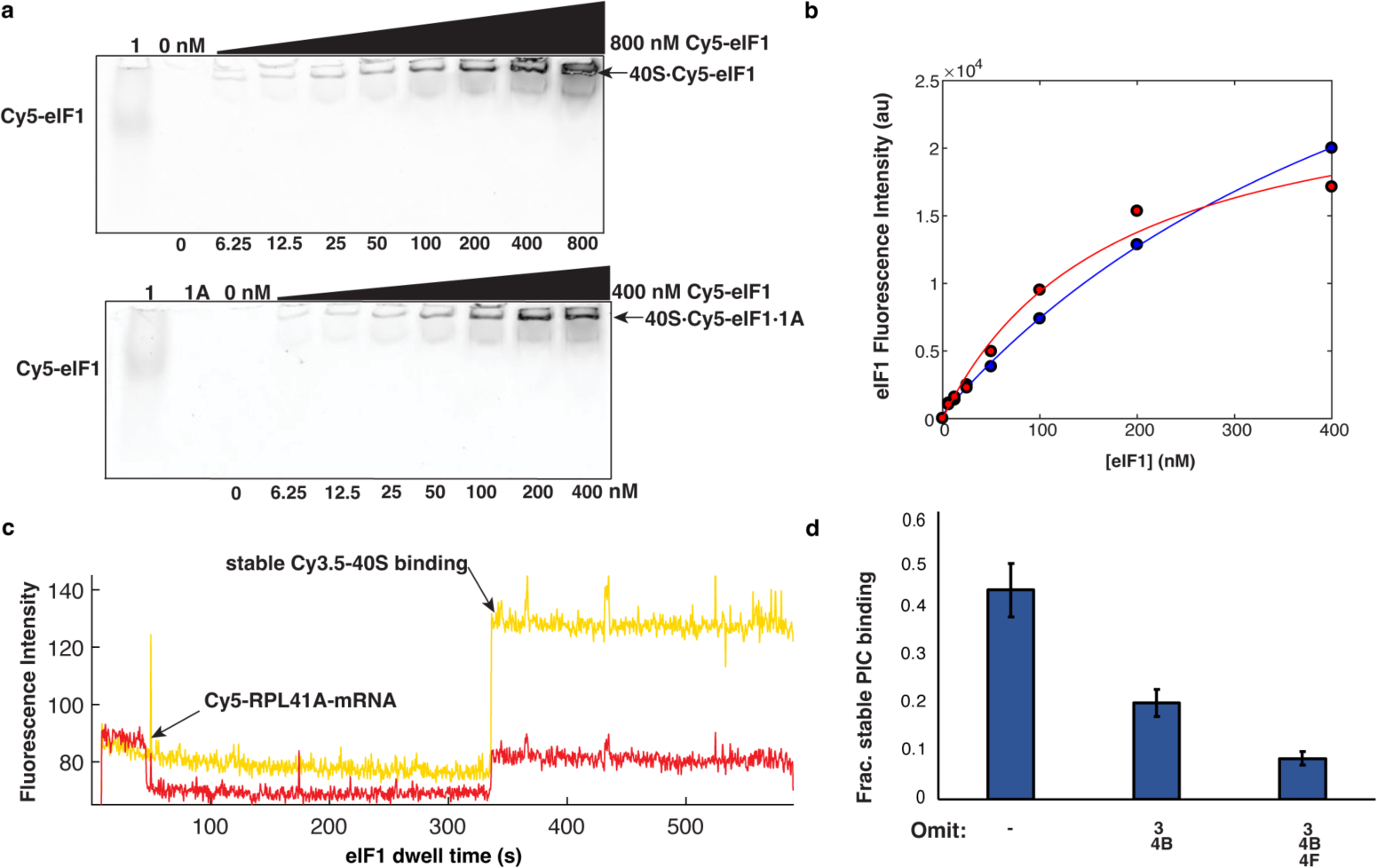
Validation of PIC assembly and on-pathway PIC–mRNA loading. **a.** Gel shift assay of Cy5-labelled eIF1 titrated against 40S subunits in absence and presence of eIF1a. **b.** 40S-associated eIF1 band intensities were fit to the Langmuir isotherm (K_D_ = 575.7 without eIF1A (blue), and K_D_= 175.7 with eIF1A (red)) **c.** Idealised single-molecule fluorescence trace for Cy5-labelled *RPL41A* mRNA with Cy3.5-40S stably binding to the RNA. The experiment contains eIFs 1, 1A, 3, 5, 4A, 4G, 4B, eIF2 ternary complex, along with ATP and GTP. **d.** Fraction of stable PIC binding under the assay conditions from panel c, modified by omission of the indicated factor(s). Error bars represent the standard deviation from 10,000 bootstrap samples of four randomly chosen sets of molecules within each condition (*n* = 148, 144, 123, 142 molecules for the full PIC; 137, 735, 423, 377 for –eIF3/4B; 1,255, 1,182, 447, 391 for –eIF3/4B/4F, respectively).

**Extended Data Figure 3.**
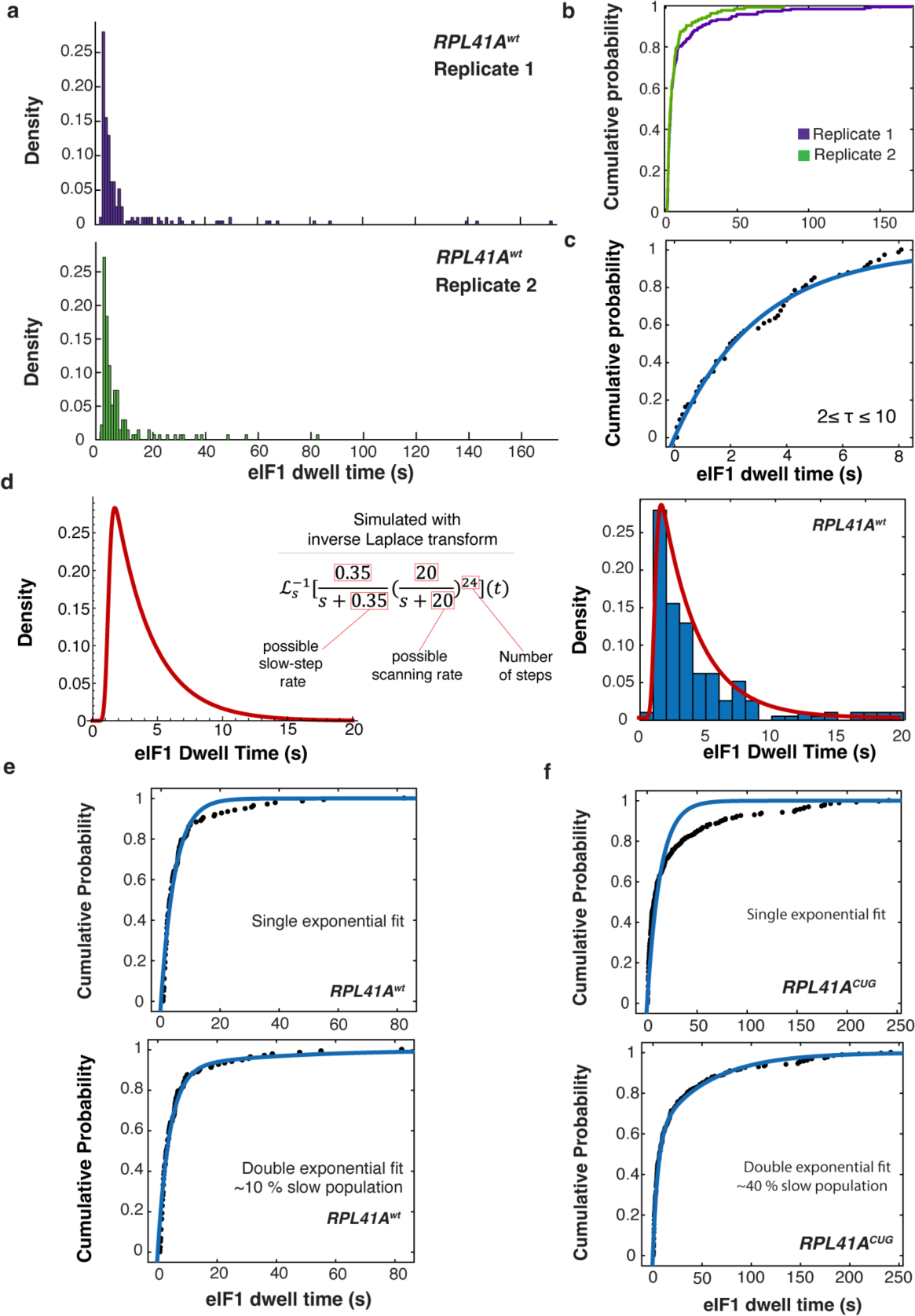
Evaluation of data reproducibility and kinetic modeling. **a.** eIF1 dwell-time distribution for the *RPL41A* mRNA in two replicates (*n* = 193 molecules for the first replicate and *n* = 136 molecules for the second replicate). **b.** eIF1 dwell-time cumulative distribution functions for both replicates of RPL41A^wt^. *p*-value was calculated using Wilcoxon rank-sum test, to evaluate the equality of the two distributions (*p* = 0.46). **c.** Exponential fit to dwell-time data extracted from the 2 – 10 s time domain of the *RPL41A*^wt^ distribution. **d.** eIF1 dwell time distribution simulation with inverse Laplace transform for *RPL41A*^wt^. **e.** eIF1 dwell-time cumulative distribution functions for scanning on *RPL41A*^wt^ fitted to a single (top) and double exponential fit (bottom). There is approximately 10 % slow population for *RPL41A^wt^*. **f.** eIF1 dwell-time cumulative distribution functions for scanning on *RPL41A*^CUG^ fitted to a single (top) and double exponential fit (bottom). There is approximately 40 % slow population for *RPL41A^CUG^*.

**Extended Data Figure 4.**
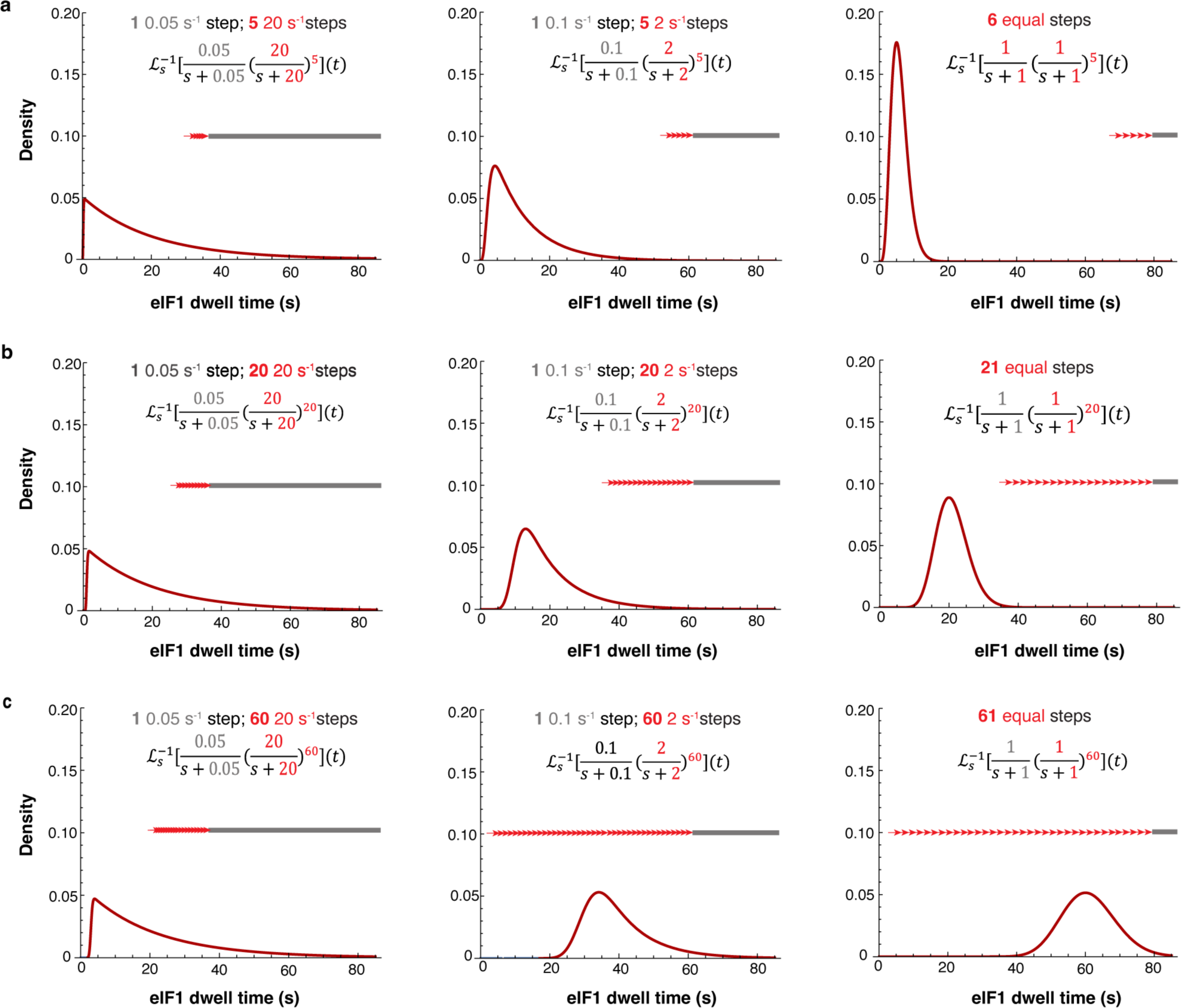
Simulation of scanning as a multi-step kinetic process. Simulated probability density functions for eIF1 dwell-time distributions under kinetic regimes with varying numbers (top to bottom) and relative timing (left to right) of multiple fast and a single slow step. The model assumes all steps are irreversible. Distributions are generated by inverse Laplace transforms, corresponding to convolution of multiple real-time (exponential) probability density functions. When the single slow step dominates the total passage time, then the distribution is dominated by a single decaying exponential component. However, as the multiple fast steps increasingly dominate the passage time, the distribution becomes more peaked, approaching a normal distribution. **a.** Models for a pathway of six sequential steps where the multiple fast steps occupy an increasing proportion of the total passage time, moving from left to right. **b.** Models for a 21-step pathway where an increasing proportion of the total passage time is dominated by multiple fast processes. **c.** Models for a 61-step pathway where an increasing proportion of the total passage time is dominated by multiple fast processes. The distribution envelopes move further from *t* = 0 as the number of steps increases, and become narrower with respect to their mean values. The schemes are depicted, for comparison with scanning, with the slow step shown at the end of the sequence. However, equivalent distributions will be obtained regardless of the relative ordering of fast and slow steps.

**Extended Data Figure 5.**
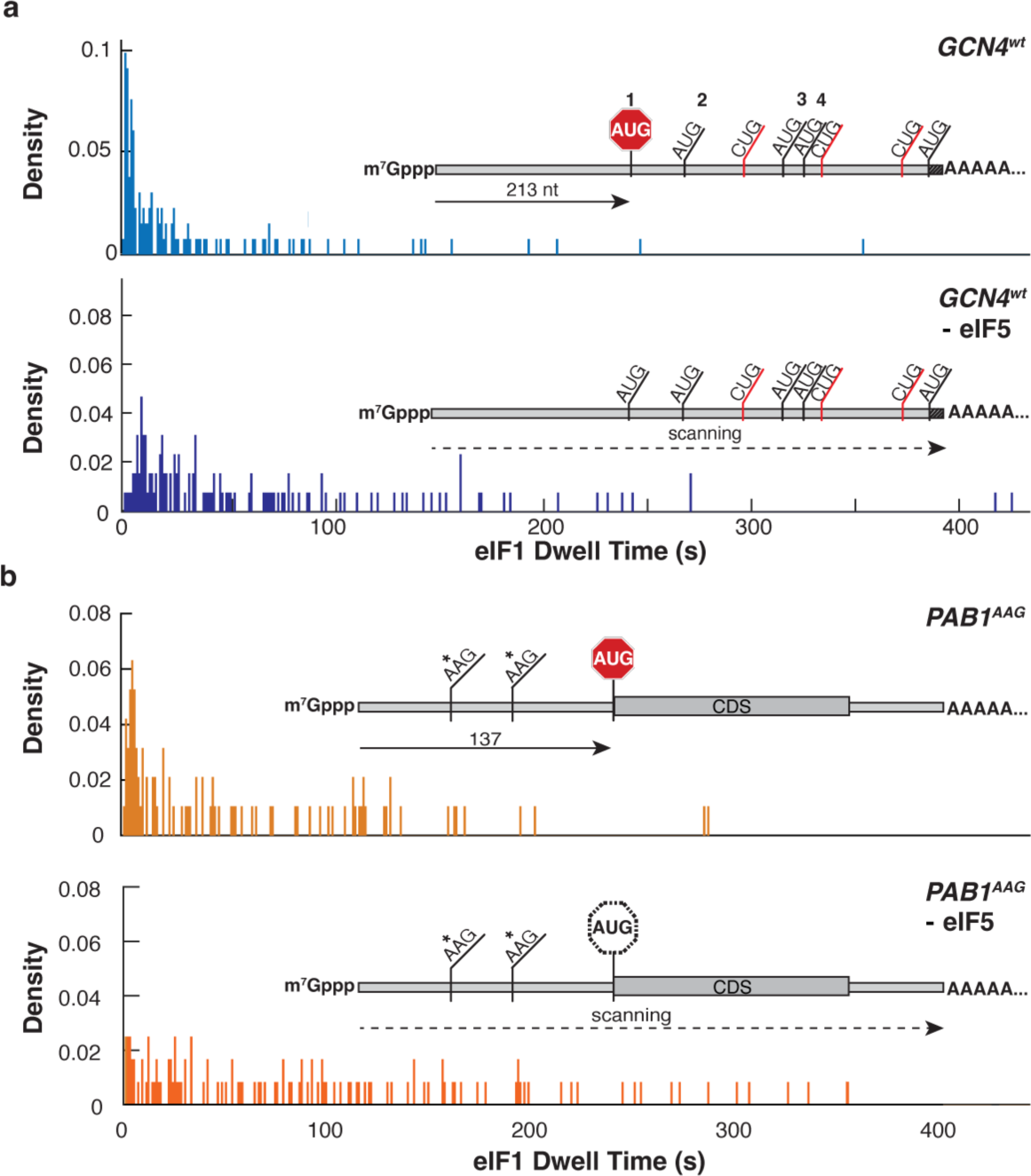
Effects of eIF5 omission on scanning dynamics. **a.** eIF1 dwell-time distribution for the *GCN4^wt^* mRNA in the presence and absence of eIF5. (*n* = 273 molecules for full PIC condition and *n* = 129 molecules for –eIF5 condition). **b.** eIF1 dwell-time distribution for the *PAB^AAG^* mRNA with and without eIF5. (*n* = 95 molecules for the full PIC condition and *n* = 122 molecules for the –eIF5 condition).

**Extended Data Figure 6.**
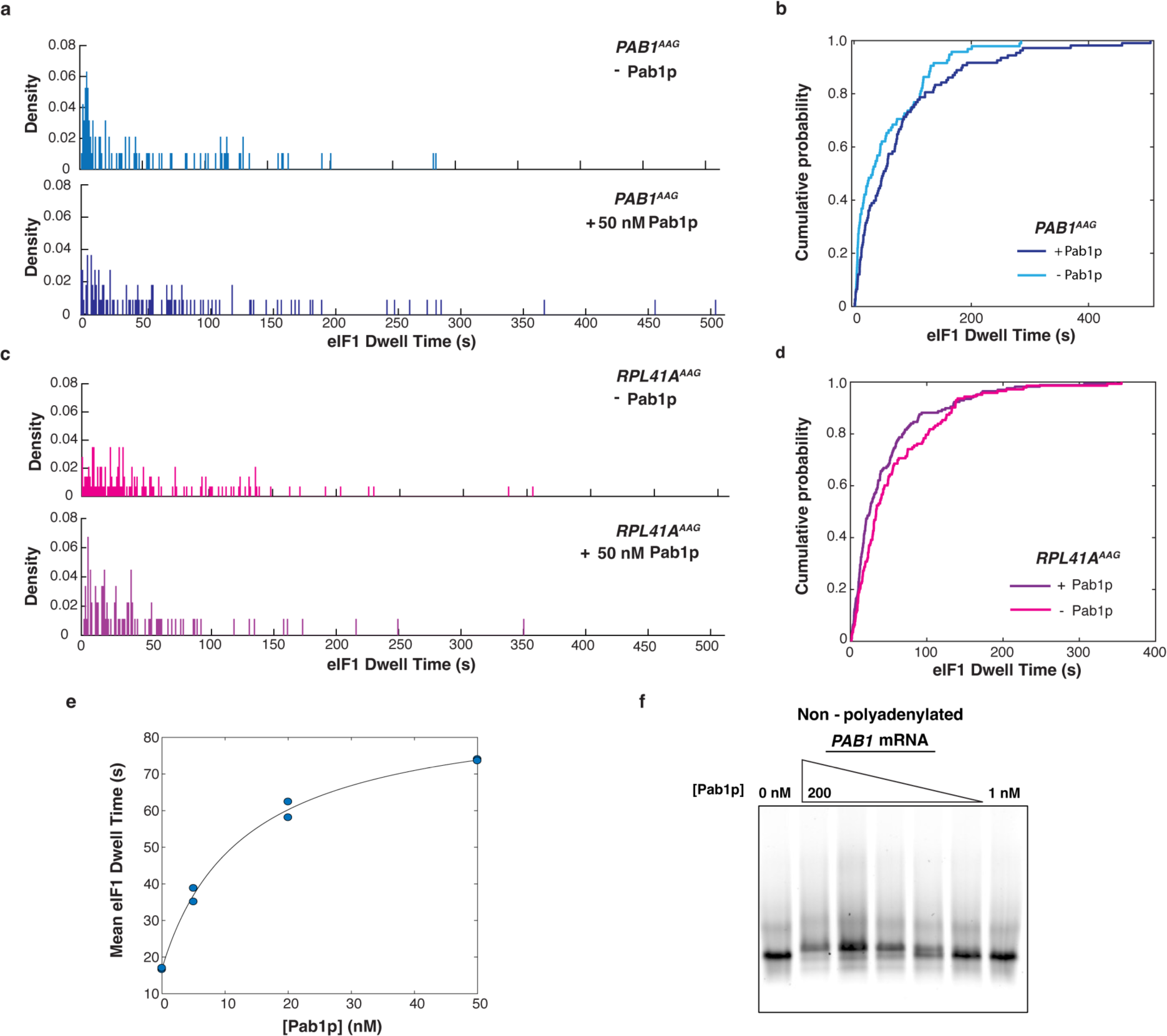
Effects of Pab1p on scanning dynamics for *PAB1*^AAG^ and *RPL41A*^wt^ mRNAs. **a.** eIF1 dwell-time distribution for the *PAB^AAG^* mRNA with and without Pab1p (50 nM). (*n* = 95 molecules in the absence Pab1p (-Pab1p); *n* = 108 molecules in the presence of Pab1p (+ 50 nM Pab1p)). **b.** eIF1 dwell-time cumulative distribution for *PAB^AAG^* with and without Pab1p (50 nM) **c.** eIF1 dwell-time distribution for the *RPL41A^AAG^*mRNA with and without Pab1p (50 nM). (*n* = 143 molecules in the absence Pab1p (-Pab1p); and *n* = 169 molecules in the presence of Pab1p (+ 50 nM Pab1p)). **d.** eIF1 dwell-time cumulative distribution functions for scanning on *RPL41A^AAG^* with and without Pab1p. **e.** Hill-Langmuir plot for dependence of mean eIF1 dwell times on the *PAB1* mRNA on the concentration of Pab1p (*n*_H_ = 0.98; 95 % confidence intervals: 0.46, 1.51). Replicate data points at 0 nM and 50 nM Pab1p overlap. **f.** Native electrophoretic mobility shift assay for non-polyadenylated *PAB1* mRNA binding, with varying concentrations of Pab1p (200 to 1 nM) respectively.

**Extended Data Figure 7.**
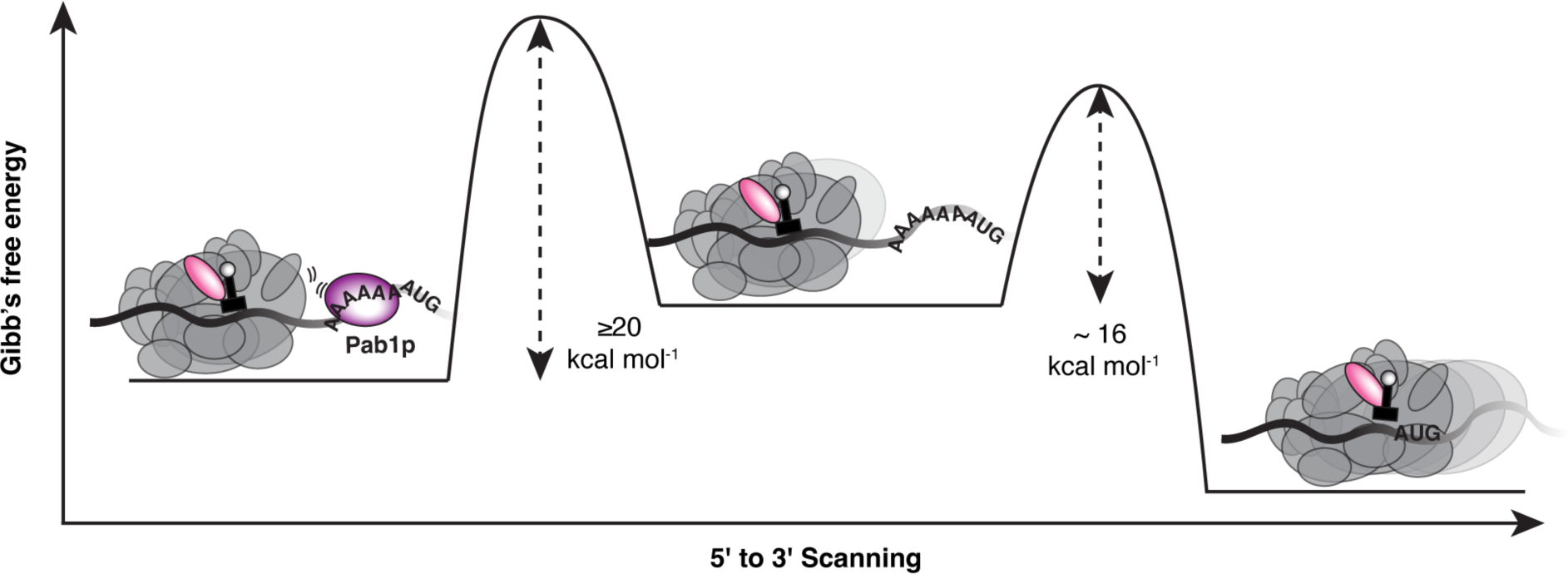
Mechanistic model for *trans-*regulation of *PAB1* scanning by Pab1p. A steric block to scanning is established by reversible Pab1p binding to the *PAB1* mRNA leader. The proportion of blocked leaders in a population of *PAB1* mRNAs rises saturably with Pab1p concentration. When the PIC arrives at the blockage, it is obliged to wait for Pab1p dissociation before it may resume normal scanning. The scanning delay time added by Pab1p corresponds to a barrier of ∼20 kcal mol^−1^ for Pab1p–*PAB1* dissociation.

